# Pulling the strings of cell cycle: a non-coding RNA, CcnA, modulates the master regulators CtrA and GcrA in *Caulobacter crescentus*

**DOI:** 10.1101/756452

**Authors:** Wanassa Beroual, Karine Prévost, David Lalaouna, Nadia Ben Zaina, Odile Valette, Yann Denis, Meriem Djendli, Gaël Brasseur, Matteo Brilli, Robledo Garrido Marta, Jimenez-Zurdo Jose-Ignacio, Eric Massé, Emanuele G. Biondi

## Abstract

Bacteria are powerful models for understanding how cells divide and accomplish global regulatory programs. In *Caulobacter crescentus*, a cascade of essential master regulators supervises the correct and sequential activation of DNA replication, cell division and development of different cell types. Among them, the response regulator CtrA plays a crucial role coordinating all those functions. Here, for the first time we describe the role of a novel factor named CcnA, a cell cycle regulated ncRNA located at the origin of replication, presumably activated by CtrA and responsible for the accumulation of CtrA itself. In addition, CcnA may be also involved in the inhibition of translation of the S-phase regulator, GcrA, by interacting with its 5’ untranslated region (5’-UTR). Performing *in vitro* experiments and mutagenesis, we propose a mechanism of action of CcnA based on liberation (*ctrA*) or sequestration (*gcrA*) of their ribosome-binding site (RBS). Finally, its role may be conserved in other alphaproteobacterial species, such as *Sinorhizobium meliloti*, representing indeed a potentially conserved process modulating cell cycle in *Caulobacterales* and *Rhizobiales*.

## Introduction

*Caulobacter crescentus* is a pivotal model organism to understand how basic functions of the cell physiology are organized and coordinated through the cell cycle (Collier, 2012; Skerker and Laub, 2004) (Figure 1A). *C. crescentus* combines the cultivation and genetic simplicity of a prokaryotic system with a regulatory intricacy that is a paradigm of global regulatory programs of all living organisms.

**Figure 1.**
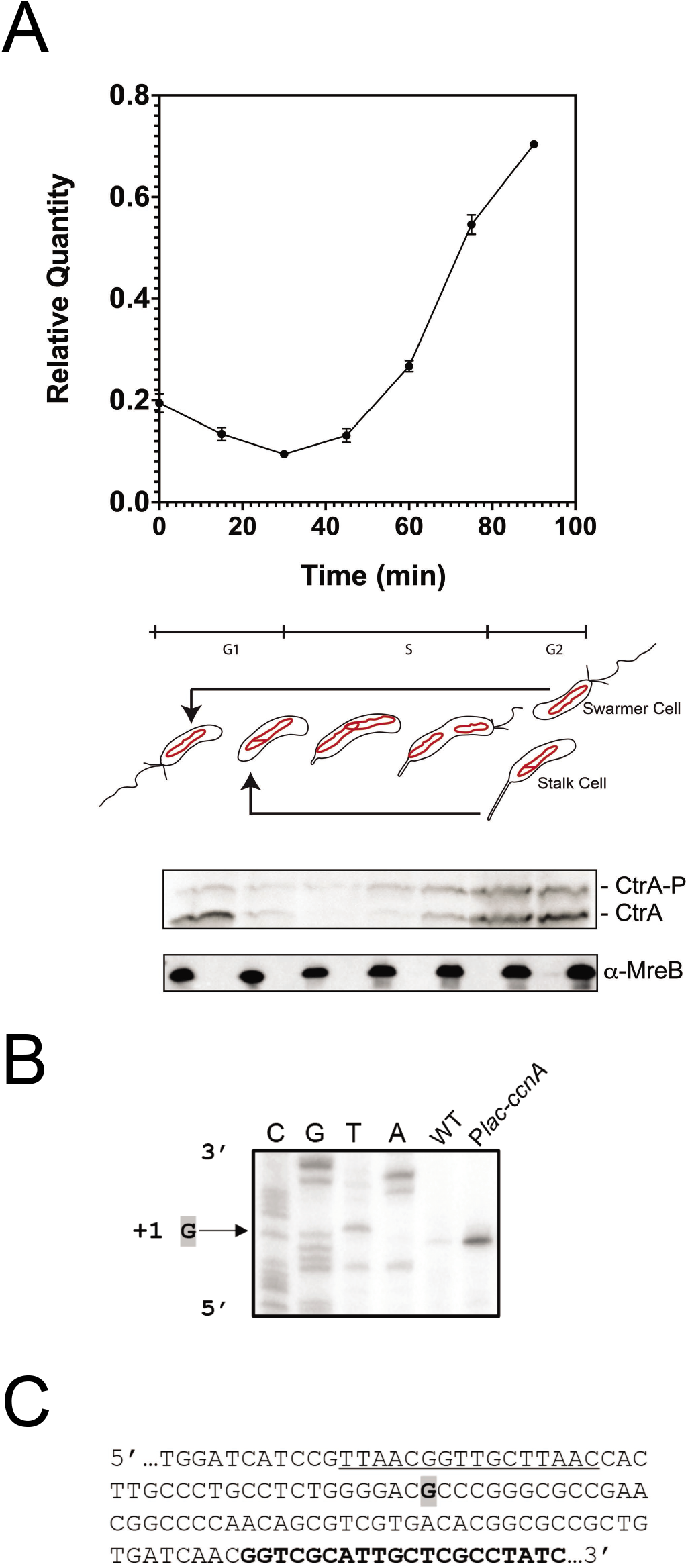
CcnA (Cell cycle Non-coding RNA A) is a cell cycle regulated ncRNA. **A. Expression level of CcnA during the cell cycle of wild-type cells *C. crescentus* (WT).** Cells were grown in PYE until OD_600nm_ = 0.6 then synchronized according to material and methods. Total RNA was extracted at indicated time points of the cell cycle. Expression of CcnA was then determined by qRT-PCR in comparison to 16S rRNA level. Results are shown as mean (N=3) +/- SD. Proteins corresponding to the same time points were extracted and separated on a SDS-PAGE gel containing Phostag and Mn^2+^ to visualize CtrA phosphorylation. CtrA (Phostag) and MreB (in a normal western blot) were revealed using specific polyclonal antibodies on nitrocellulose membranes. **B. Determination of the transcriptional +1 site of CcnA ncRNA by primer extension**. Total RNA extracted from WT cells or containing P*lac-ccnA* was used with a radiolabelled oligo (bold sequence in **C**). The same oligo was used for *ccnA* sequencing (CGTA). The sequence is presented as the reverse complement. The +1 signal is represented by the arrow. See supplementary Figure S2A for controls. Data are representative of two independent experiments. **C. DNA 5’sequence of *ccnA*.** Boxed gray “**G”** corresponds to the transcriptional +1. Oligo use for the sequencing and primer extension is in bold. CtrA box promoter region is underlined.

Transcriptional regulation plays a major role during cell cycle progression. Several master regulators controlling transcription (i.e. DnaA, GcrA, CcrM and CtrA) are sequentially activated in order to induce transcription of hundreds of genes required at specific phases of the cycle (Collier et al., 2006, 2007; Reisenauer and Shapiro, 2002). Each phase is under the control of well-defined factors: (i) the initiation of the S-phase depends on DnaA, (ii) the first part of the S-phase depends on the epigenetic module GcrA and CcrM and (iii) the second part depends on CtrA, which is also the regulator of the G1 phase of swarmer cells (Panis et al., 2015).

Other regulators of transcription intervene to fine tune the cell cycle-regulated transcription of genes that must be activated at specific phases of the cell cycle; for example, MucR and SciP regulate CtrA activity (Delaby et al., 2019; Fumeaux et al., 2014; Gora et al., 2010, 2010, 2013). The interconnections between DnaA, GcrA, CcrM and CtrA create an intricate network whose behavior emerges from the integration of multiple master regulatory inputs. In particular, regulation of the essential response regulator CtrA is critical, as it directly or indirectly controls all the other master regulators of the cell cycle (Laub et al., 2002). CtrA is notably responsible for the direct transcriptional activation of key genes for cell division and the biogenesis of polar structures (flagellum, stalk and pili). CtrA also activates the transcription of the gene encoding the orphan adenine methyl transferase CcrM, which in turn is required for the regulation of many genes including the fine-tuned regulation of the promoter P1 of *ctrA* (Reisenauer and Shapiro, 2002). Moreover, CtrA indirectly blocks chromosome replication initiation promoted by DnaA by binding to sites in the origin of replication (*CORI*), resulting in DnaA exclusion from the *CORI* (Marczynski and Shapiro, 2002; Quon et al., 1998).

Another master regulator, named GcrA, activates the transcription of the *ctrA* gene, which in turn negatively feeds back on the transcription of *gcrA* (Fioravanti et al., 2013; Haakonsen et al., 2015a; Holtzendorff et al., 2004; Mohapatra et al., 2020). GcrA activity is known to be affected by the methylation status of its targets’ promoters. For instance, the GcrA–dependent transcription of *ctrA* from its P1 promoter is activated by the conversion of a CcrM methylated site from its full to the hemi-methylation state approximately after a third of DNA replication (Reisenauer and Shapiro, 2002). P1 activation is therefore responsible for the first weak accumulation of CtrA, and predates the activation of the stronger P2 promoter, located downstream of P1. P2 is under the control of phosphorylated CtrA (CtrA~P), responsible for the robust accumulation of CtrA in the second half of DNA replication. CtrA at its highest level is then responsible for the repression of its own P1 promoter and of *gcrA* transcription. Although the molecular details of this biphasic activation of *ctrA* are still only partially understood, the stronger activation of P2 may underscore other post-transcriptional mechanisms reinforcing CtrA accumulation.

Besides being finely regulated in time by the DnaA-GcrA-CcrM transcriptional cascade, activation of CtrA requires phosphorylation by the CckA-ChpT phosphorelay (Biondi et al., 2006a), which is linked to a sophisticated spatial regulation since the hybrid kinase CckA has a bipolar localization (Biondi et al., 2006a; Chen et al., 2011; Jacobs et al., 2003). At the swarmer pole, CckA acts as a kinase due to the presence of the kinase DivL and the DivK phosphatase PleC (Gora et al., 2010). However, at the stalked pole, CckA is a phosphatase of CtrA because the kinase DivJ keeps the CtrA negative regulator DivK fully phosphorylated, turning the CckA-ChpT phosphorelay into a CtrA phosphatase. As CtrA~P blocks the origin of replication, a complex degradation machinery ensures its cell cycle-dependent degradation at the G1 to S-phase transition and after cell division in the stalk compartment. A cascade of adapter proteins (CpdR, RcdA and PopA) is responsible for the specific and highly regulated proteolysis of CtrA (Joshi et al., 2015; Ryan et al., 2004).

Few cases of regulation of gene expression by ncRNAs have been characterized in *C. crescentus*. For example, the SsrA non-coding RNA (tmRNA) is a small RNA associated to selected translating ribosomes to target the translated polypeptides for degradation. tmRNA has been linked to replication control in *C. crescentus* (Keiler and Shapiro, 2003) and *Escherichia coli* (Wurihan et al., 2016). More generally, only 27 ncRNAs were described in *C. crescentus* (Landt et al., 2008). Among them, CrfA is a ncRNA involved in adaptation to carbon starvation (Landt et al., 2010). Another ncRNA, GsrN, is involved in the response to multiple σ^T^-dependent stresses (Tien et al., 2018). Finally ChvR has been recently characterized as a ncRNA that is expressed in response to DNA damage, low pH, and growth in minimal medium (Fröhlich et al., 2018). However, as more recent approaches using RNA sequencing (RNAseq) and post-genomic techniques expanded the *plethora* of ncRNA candidates to more than 100 (Zhou et al., 2015). Predictions of their integration into the cell cycle circuit (Beroual et al., 2018) suggest that those new candidate ncRNAs should be deeply studied in order to find whether ncRNAs are linked to cell cycle regulation. Indeed, ncRNA-mediated regulations can provide network properties that are not always easily accessible through transcriptional regulation only. For example, the phenomena like threshold-linear response of the mRNA target, the prioritization of different targets, ultrasensitive response and bistability are known regulatory mechanisms mediated by ncRNAs (Levine et al., 2007; Mitarai et al., 2009), therefore representing good candidates as regulators of a biological system showing rich dynamic behavior as the *C. crescentus* cell cycle.

Here we investigated the role of a ncRNA, named CcnA, which is transcribed from a gene located at the origin of replication of the *C. crescentus* chromosome. We characterized its role in cell cycle regulation by using deletion mutants, CcnA overexpression strains and silenced strains obtained through expression of a CcnA antisense RNA. Results presented here identified the mRNAs of CtrA and GcrA, two master regulators of cell cycle, as important targets of the CcnA ncRNA. Our results are supported by a multipronged approach, combining “MS2-affinity purification coupled with RNA sequencing” (MAPS) assays, *in vitro* and *in vivo* experiments. Finally, the role of CcnA in the closely related organism *Sinorhizobium meliloti* suggests an evolutionary conservation across alphaproteobacteria, further underscoring the importance of this gene.

## Results

### CcnA expression is activated in predisional cells

Based on previous results (Zhou et al., 2015), we speculated that CCNA_R0094, here named <u>C</u>ell <u>C</u>ycle <u>n</u>on-coding RNA <u>A</u> (CcnA), has its peak of transcription after the accumulation of CtrA, in the second half of the S-phase, when, the second *ctrA* promoter, P2, is activated. A synchronized population of wild type *C. crescentus* was used to collect cells at 15 minutes intervals in rich medium (generation time is 96 minutes). We designed primers to detect and precisely quantify CcnA RNA in the cells during cell cycle by q-RT-PCR (see Materials and methods) with respect to 16S RNA levels (Figure 1A). CcnA levels start increasing after 45 minutes, coincidentally with CtrA protein levels (Figure 1A). More specifically, we measured both protein and phosphorylation levels of CtrA by Phos-Tag gels (Figure 1A). CcnA levels increase as CtrA~P levels increase, suggesting that the transcription of *ccnA* potentially depends on phosphorylated CtrA. This observation prompted us to question whether CtrA was involved in *ccnA* transcription. Consistent with this, a CtrA box was previously described upstream of the Transcriptional Start Site (TSS) of *ccnA* (Brilli et al., 2010a; Zhou et al., 2015).

We performed RNA-seq using a *ctrA* thermo-sensitive allele *ctrA401ts* (*ctrA*-ts) to test for variations of *ccnA* expression in the context of the global transcriptional changes taking place in this highly perturbed mutant (Biondi et al., 2006a; Laub et al., 2002; Quon et al., 1996). At the permissive temperature (30**°C)**, *ctrA-ts* shows a partial loss of function phenotype while the strain doesn’t grow at the restrictive temperature (37**°C)** (Quon et al., 1996). The analysis on *ccnA* revealed that expression of CcnA is reduced in the *ctrA*-ts compared to wild type at the restrictive temperature (Figure S1A), suggesting that of completely functional CtrA is required to express *ccnA*. This observation is consistent with the cell cycle regulated profile of CcnA and with a predicted CtrA binding site in the *ccnA* promoter region. This result was further supported (Figure S1B) by the observation that CcnA shows increased levels in strains where CtrA has higher levels of stability, such as *rcdA, popA* and *cpdR* mutants in which CtrA protein steady state levels are higher than the WT (Figure S1C). The increase of *ccnA* transcription in *rcdA, popA, cpdR* and *divJ* deletion strains indeed support the hypothesis that *ccnA* transcription may be regulated by CtrA.

In summary, CcnA is a ncRNA regulated by cell cycle and putatively regulated in a positive way by CtrA, with peak expression in the second half of DNA replication, coincident with CtrA accumulation. Considering the high affinity of CtrA on the promoter of CcnA (Siam et al., 2003), future studies are necessary in order to investigate this putative CcnA transcriptional activation by CtrA.

### CcnA transcription is required for the accumulation of CtrA

To understand the function of CcnA by overexpression, we fused the sequence of *ccnA* with the first transcribed nucleotide of a P*lac* promoter in the vector pSRK (Khan et al., 2008) (see Materials and Methods). This vector was introduced in *C. crescentus* cells and its +1 nucleotide was analyzed in the over expression strain in comparison with the wild type native CcnA by primer extension (Figure 1, B, C) (see Materials and Methods). The level of CcnA in this inducible system, estimated by primer extension (Figure 1B, Figure S2A) and quantified by q-RT-PCR (Figure S1B), confirmed higher levels of CcnA expression compared to the wild type. Cells overexpressing *ccnA* showed cell cycle defects, such as slow growth (Figure 2A, B, C), morphologies related to abnormal cell division (Figure 2D), with an increased number of long stalks (Figure S3). Several tests were performed in order to characterize these phenotypes and provide a basis to better understand the mechanisms behind them. By quantifying cell size parameters by using MicrobeJ (Ducret et al., 2016a), we discovered that cells expressing *ccnA* ectopically were significantly more elongated and filamentous than wild type cells (Figure 2E). Stalk biogenesis, cell division and inhibition of DNA replication are all under the control of CtrA (Biondi et al., 2006b; Quon et al., 1996) suggesting that CcnA may feedback on CtrA production to affect these processes. Indeed, upon expression/overexpression of CcnA, CtrA accumulates to higher steady state levels with respect to the control strain, while in the loading control, MreB, levels are constant (Figure 2F). We also checked the effect of high CtrA levels on the DNA replication behavior. As previously demonstrated, the over expression of CtrA in a WT background does not induce a block of DNA replication given its cell cycle regulated proteolysis (McGrath et al., 2006). Flow cytometry experiments showed that over expressing CcnA indeed did not induce a change in DNA content (Figure S4A, B).

**Figure 2.**
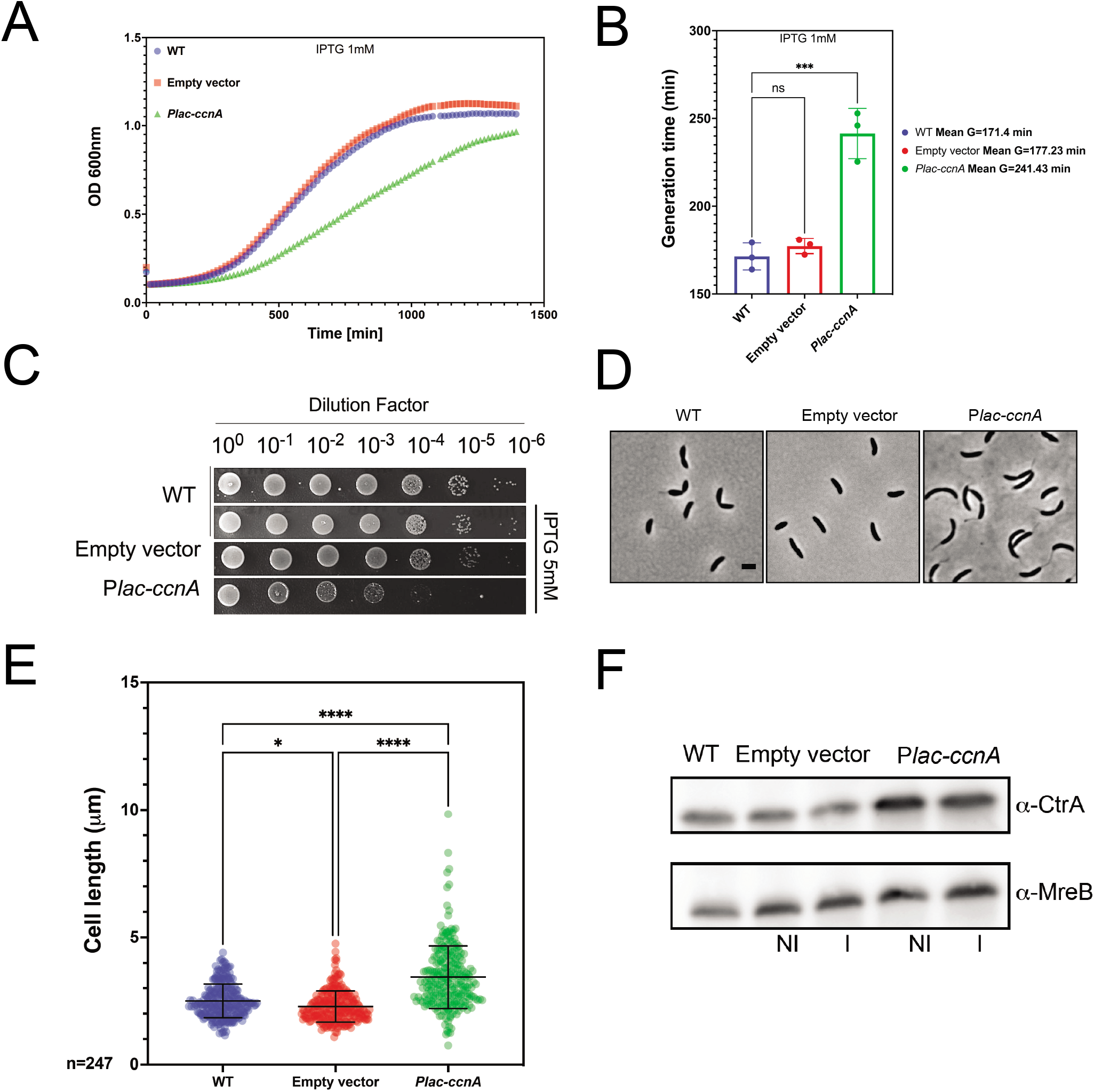
CcnA affects the cell cycle. **A. Growth curves following the expression of CcnA.** WT cells and WT cells carrying either an empty pSRK (empty vector) or a pSRK with *ccnA* under the control of an inducible P*lac* promoter (P*lac-ccnA*) were grown in PYE without IPTG. 200μL of cells back-diluted from stationary phase cultures to an OD_600nm_= 0.02 were then grown on 96 wells in PYE supplemented with 1mM IPTG. Cell growth was monitored overnight with a Spark-TM at 30°C and a Shaking (orbital) Amplitude of 6 mm and a shaking (orbital) frequency of 96 rpm. Results are shown as mean N=3 biological replicates with 3 technical replicates. Raw data are provided in Table S7. **B. Determination of the doubling time of cells expressing CcnA**. Doubling times of cells from (2A) were calculated by using the exponential growth equation (Nonlinear regression) (Prism GraphPad 9.1.2). Statistical analysis was performed using ANOVA with a Brown-Forsythe and Welch ANOVA tests and a Dunnett’s multiple comparisons test. ns= difference not significant, ***: p.val = 0,0002. **C.** WT cells, WT cells carrying either an empty vector or P*lac-ccnA* were grown overnight in PYE at 30°C and diluted to an OD_600nm_= 0.6. Samples were then serially diluted (10^0^–10^−6^) and 4.5μL of each dilution were spotted on a PYE-Agar + 5mM IPTG plate and incubated at 30°C. WT cells without plasmid were used as negative control. **D.** Phase contrast images of WT cells, WT cells carrying an empty vector or P*lac-ccnA* grown in PYE without IPTG until OD_600nm_=0.6. Scale bar = 2μm. **E.** Cells from (2D) were analyzed using MicrobeJ (Ducret et al., 2016a) to assess cell length. 247 cells were analyzed for each condition and statistical significance was determined using ANOVA with Tukey’s multiple comparisons test. *: p.val=0.0168 ****: p.val<0.0001. Raw Data are provided in Table S8. **F.** WT cells, WT cells carrying an empty vector or *Plac-ccnA* were grown in PYE at 30°C until OD_600nm_ = 0.6. Then, induction of P*lac-ccnA* was made by addition of IPTG 1mM 30min. As a control of induction WT cells carrying an empty vector were also incubated 30min in presence of IPTG 1mM and WT cells with no induction were used as a control (NI= no IPTG) and (I= IPTG). Proteins were extracted and separated on a SDS-PAGE gel for Western blotting. CtrA and MreB (loading control) proteins were revealed using specific polyclonal antibodies on nitrocellulose membranes. Results are representative of at least three independent experiments (See Figure S14 for additional Westerns).

CtrA must be phosphorylated by a phosphorelay that includes CckA and ChpT to become fully active (Figure S5A). The Phos-tag technique, implemented as previously described (Pini et al., 2013) allowed us to evaluate the levels of CtrA~P upon overexpression of CcnA. The analysis revealed that the band of CtrA~P was more intense than the band of non-phosphorylated CtrA when CcnA was overexpressed (Figure S5B). As phosphorylation of CtrA is under the control of the phosphorelay CckA-ChpT (Biondi et al., 2006a), we tested whether levels of ChpT were also affected, which would indicate an increased activity of the phosphorelay. We used a YFP translational fusion of ChpT (ChpT-YFP) in order to understand whether CcnA ectopic expression was causing a change in protein subcellular localization and levels. Epifluorescent microscopy was used to observe the protein level of ChpT-YFP (Figure S5C). Data were further analyzed by MicrobeJ (Materials and Methods) and results were compared to a strain carrying an empty vector showing that upon CcnA overexpression intensity and clustering of the signal increase in the ChpT-YFP strain background, more specifically in elongated cells with long stalks (Figure S5D, E). Finally, we tested by Western-blot whether CcnA overexpression affected the protein level of ChpT, using antibodies against the GFP protein that does recognize YFP and compared the levels of the ChpT-YFP translational fusion in strains carrying either an empty vector or CcnA. Our results showed that upon overexpression of CcnA, YFP-ChpT levels were higher than those of the empty vector (Figure S5F). This observation may suggest that CcnA overexpression increases CtrA phosphorylation by affecting the localization and levels of ChpT by an unknown mechanism so far.

In conclusion, an increase in CcnA expression induces an increase in the steady state levels of CtrA protein, specifically in its phosphorylated form (CtrA~P). These changes in the CtrA levels may well explain the cell cycle defects observed at the morphological and molecular levels, notably increase of cell length, and long stalks.

The gene *ccnA* is located in the origin of replication (Figure 3A); therefore, its sequence, at least partially, plays an essential role in the initiation of replication (Christen et al., 2011a; Taylor et al., 2011). We attempted a complete deletion of the *ccnA* sequence by two-step recombination in the presence of an extra copy of *ccnA* (Materials and Methods), as previously described (Skerker et al., 2005). Considering that *ccnA* coincides with an essential part of the origin of replication of the genome, the deletion of the *ccnA* sequence was not successful, demonstrating that the genomic sequence of *ccnA* is essential (Christen et al., 2011a). We then applied different strategies to inactivate partial sequences of *ccnA* that kept most of the origin of replication intact (Figure S6) without success. Finally, we attempted to delete the 45 bp long promoter region containing the CtrA box. The *ccnA* expression should be under the control of CtrA, therefore we hypothesized that the deletion of its box in the promoter region should have a mild or no effect on the origin but impair the expression of the ncRNA. The deletion of the promoter region was obtained and the expression of *ccnA* in the corresponding mutant was first tested by primer extension (Figure 3B, Figure S7A) that showed the absence of CcnA. We also used qRT-PCR (Figure S1B) using primers for *ccnA* and the 16S sequence as reference (Materials and Methods) in order to quantify the decrease of CcnA upon deletion of its putative promoter (Δ*prom* mutant). Upon deletion of the promoter region we observed a significant decrease of CcnA expression that may explain the cell cycle defects (phenotypes that are similar to silencing approach, see below) (Figure S1B).

**Figure 3.**
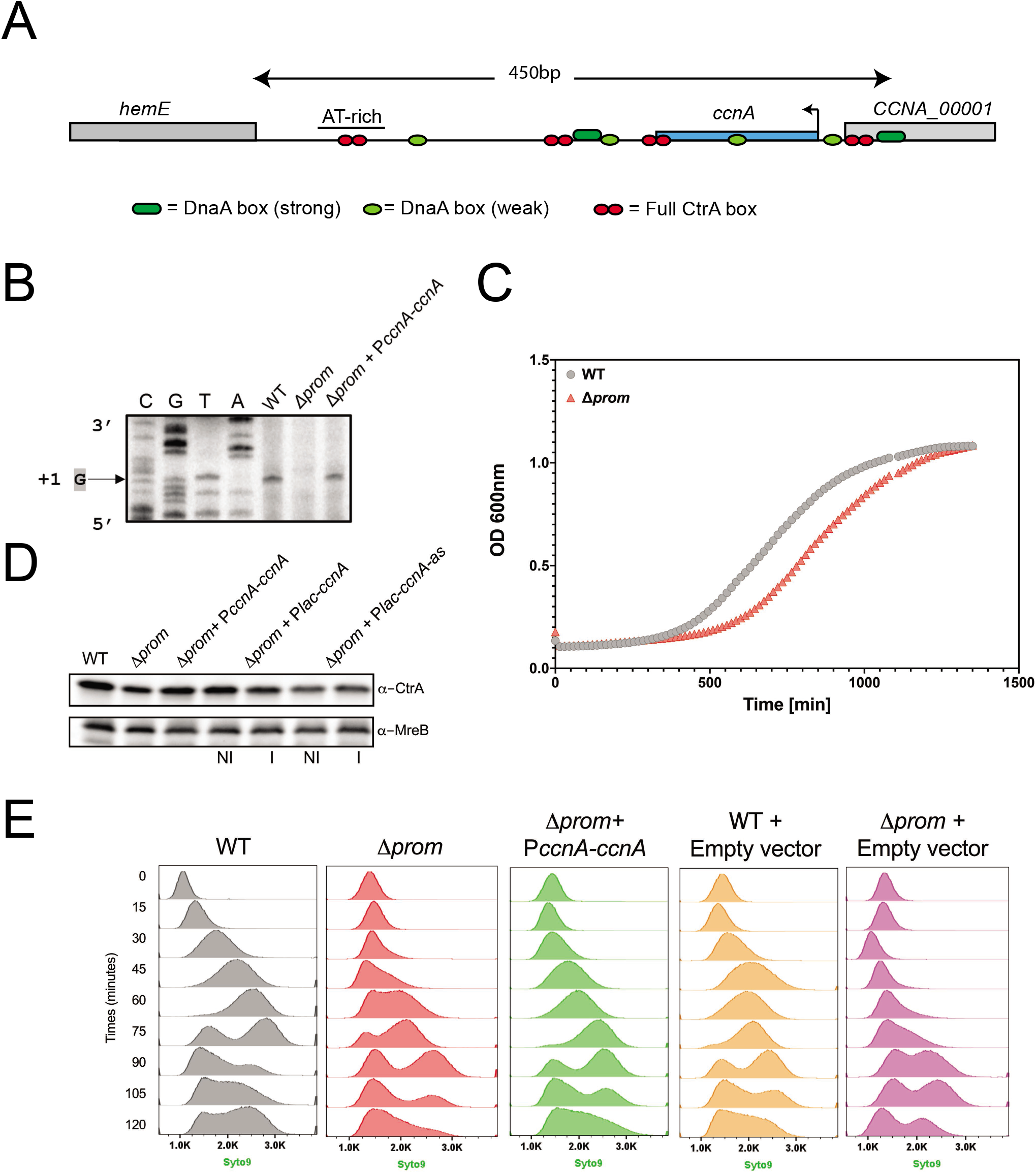
Δ*prom* cell cycle defects are rescued by CcnA *in trans* under the control of its own promoter. **A. Schematic representation of the origin of replication and *ccnA* gene locus in *C. crescentus***. The origin of replication contains 4 full CtrA boxes, 2 strong and 4 weak DnaA boxes (Frandi and Collier, 2019). Transcription of *hemE* gene serves as nucleotides template for DNA replication (Okazaki fragments) (Marczynski et al., 1995). The chromosome replication initiator DnaA unwinds the DNA from the AT Rich region on the chromosome when CtrA is absent. The *ccnA* gene is 182 nt-long and contains a CtrA box in its promoter region and in its terminal region. **B. Determination of the transcriptional +1 site of CcnA ncRNA by primer extension**. Total RNA extracted from wild-type cells (WT), deleted *ccnA* promoter (*Δprom*) and containing P*ccnA-ccnA* (Δ*prom* + P*ccnA-ccnA*) were used with radiolabelled oligo. The same oligo was used for *ccnA* sequencing (CGTA). The +1 signal is represented by the arrow. See supplementary figure S7A for controls. Data are representative of two independent experiments. **C. Growth curves of cells deleted from *ccnA* promoter.** WT cells and Δ*prom* cells were grown overnight in PYE at 30°C. 200μL of cells back-diluted from stationary phase cultures to an OD_600nm_= 0.02 were then grown on 96 wells in PYE. Cell growth was monitored overnight with a Spark-TM at 30°C and a Shaking (orbital) Amplitude of 6 mm and a shaking (orbital) frequency of 96 rpm. Results are shown as mean N=2 biological replicates with 3 technical replicates. Raw data are provided in Table S7. **D**. WT cells, Δ*prom* cells, Δ*prom* cells carrying either a pMR10 low copy plasmid harboring *ccnA* under the control of its own promoter (Δ*prom* + P*ccnA-ccnA*) or *ccnA* or its antisense under the control of a P*lac* promoter (Δ*prom*+ P*lac-ccnA*, Δ*prom* + P*lac-ccnA-as*) were grown in PYE at 30°C until OD_600nm_= 0.6. For Δ*prom*+ P*lac-ccnA*, Δ*prom* + P*lac-ccnA-as* cells, expression of *ccnA* or its antisense was made by addition of IPTG 1mM 30min. Proteins were extracted and separated on a SDS-PAGE gel for Western blotting. CtrA and MreB (loading control) proteins were revealed using specific polyclonal antibodies on nitrocellulose membranes. Results are representative of at least two independent experiments with similar results (See supplementary figure S7B and S14D for controls). **E.** Flow cytometry profiles after SYTO 9 staining showing DNA content of synchronized WT cells, Δ*prom* cells, Δ*prom* cells carrying *ccnA* under its own promoter (Δ*prom* + P*ccnA-ccnA*) and as controls WT cells carrying an empty low copy plasmid pMR10 (WT + empty vector) or Δ*prom* cells carrying and empty low copy plasmid pMR10 (Δ*prom* + empty vector). Synchronization of cells was performed as described in material and methods. Pure G1 (1N) swarmer cells were isolated by Percoll for density gradient and DNA replication over the cell cycle was followed on synchronized cells at different time point. A total number of 300 000 particles were analysed by flow cytometry using the blue laser (488nm) and filter 525/30 nm. Results are representative of three biological replicates.

The Δ*prom* mutant was analyzed by growth curves (Figure 3C) and its morphology was observed by microscopy (Figure S8A). This strain showed slow growth and more precisely a longer lag phase than the WT strain (Figure 3C). Western blots were performed using antibodies against CtrA and MreB (Figure 3D). This mutant showed a decrease of CtrA steady state levels, as expected considering the opposite effects in the overexpression strain (Figure 2F). On the contrary, MreB (loading control) remained stable, suggesting a specific effect on CtrA. As the deletion of *ccnA* promoter removes also some elements of the origin of replication (Taylor et al., 2011), we performed flow cytometry analysis on synchronized populations to understand whether the deletion of *ccnA* promoter of *C. crescentus* does not interfere with DNA replication initiation. Flow cytometry analysis revealed that the markerless deletion of *ccnA* promoter does not have a strong effect on DNA replication but probably causes a delay in the initiation of DNA replication (Figure 3E). Given the lower level of CtrA in the *Δprom* strain, we would expect DNA replication to occur at a higher rate in this mutant. However, we observed normal initiation of DNA replication in the WT strain with a shift in peak intensity from 30 minutes of the cell cycle demonstrating that DNA replication has begun and a total shift from 1 chromosome (1N) to 2 chromosomes (2N) at 60 minutes whereas the *Δprom* strain remained blocked with 1N until 60 minutes and began to accumulate 2N content only at 60 minutes given the second peak that was observed. We estimated the percentage of 1N in *Δprom* cells at 34.65% +/- 3.88% and 2N at 63.85+/- 3.46.

We complemented the *Δprom* strain with a WT copy of *ccnA* under the control of its own promoter in a low-copy vector (*PccnA-ccnA*). We were interested in understanding whether a deletion of a portion of *CORI* was the sole reason of the *Δprom* phenotypes or whether it was due to a lack of CcnA transcription.

Indeed, the Δ*prom* was almost entirely complemented by an extra copy of the *ccnA* gene, as DNA replication (Figure 3E), growth (Figure S8B), CtrA levels (Figure 3D) were rescued by the extra copy of CcnA, demonstrating that the phenotype of Δ*prom* was mostly related to the absence of CcnA.

An alternative, less invasive strategy with respect to the origin of replication was to overexpress an antisense of CcnA (CcnA-as) in order to silence the RNA of CcnA. A reverse complementary sequence of CcnA driven by a *Plac* promoter was cloned, as described in the previous section for the sense sequence and expressed in *C. crescentus* in order to demonstrate a negative effect on CcnA activity. Based on Western blots, the expression of the antisense of CcnA, as the Δ*prom* strain, showed a decrease of CtrA steady state levels (Figure S9A). Flow cytometry analysis also showed an accumulation of chromosomes (N≥3) in presence of CcnA antisense (Figure S9B, C). Moreover, an increase of doubling time was observed (Figure S9D). These results suggested that the expression of the antisense phenocopy Δ*prom*, so it may indicate an inactivation of CcnA activity. This result, together with the viability of the *ccnA* Δ*prom* strain, also suggests that the inactivation of CcnA is not lethal.

In conclusion, both overexpression and low levels of CcnA showed consistent results that suggested that CcnA promotes the accumulation of CtrA and possibly other genes expression products. Therefore, we wondered if this activity was due to a direct binding by CcnA to the 5’UTR of *ctrA* and potentially other genes.

### CcnA directly binds to mRNAs of *ctrA, gcrA* and other cell cycle genes

In order to identify RNAs that were targeted *in vivo* by CcnA and test whether CtrA mRNA was a direct target of CcnA, we performed the technique called MAPS (MS2-affinity purification coupled with RNA sequencing) as previously described (Lalaouna et al., 2015). This technique relies on the fusion of a ncRNA of interest with the RNA aptamer MS2 used as a tag at the 5’ of the ncRNA. MAPS approach involves the use of a protein called MS2-coat with a high affinity for the MS2 RNA aptamer. This technique allows the identification of RNAs or proteins directly interacting with a tagged RNA (Figure S10). We indeed constructed a version of CcnA tagged with an MS2 RNA aptamer able to bind the protein MS2-MBP immobilized on an amylose resin. As a negative control, an untagged *ccnA* was cloned in order to compare results specific to the MAPS technique. Strains expressing MS2-*ccnA* or *ccnA* (introduced in the same pSRK plasmid type previously used for *ccnA* overexpression) were lysed and soluble cell content was loaded onto an amylose column containing MS2-MBP fusion. RNA was purified as previously described (Materials and Methods) (Lalaouna et al., 2015).

RNAs trapped in the amylose column in presence of MS2-CcnA or non-tagged CcnA were characterized by RNAseq and results were analyzed (Materials and Methods). First, as a control, we looked for the presence of reads in the vicinity of *ccnA* only in the MS2-CcnA strains (Figure S11A). Among other candidate targets (Figure S12), the *ctrA* mRNA was detected (Figure S11B). This result is in accord with our previous results that CcnA overexpression and downregulation affected CtrA expression. The extent of CcnA-regulated targets is bigger than just *ctrA* mRNA. As shown in Figure S12 and Table S1, other mRNAs, including *gcrA* are potentially targeted by CcnA. A general observation of candidate targets of CcnA is that most of them belong to the CtrA regulon, such as those encoding motility proteins (Figure S12, Table S1).

We also tagged the 5’UTRs of *ctrA* with the MS2 aptamer (mRNAs generated by its P1 or P2 promoter) in order to determine the putative interaction with CcnA. We expressed the MS2 tagged UTRs in *C. crescentus* cells and we looked for the enrichment of CcnA in the MS2-P1 and MS2-P2 UTRs associated to the correct overexpression of the 5’UTRs (Figure S11C). We demonstrated that only the UTR of *ctrA* mRNA transcribed by the P2 promoter pulls down CcnA (Figure S11D). Although P1 obviously contains the sequence present in P2, it may form different secondary structures that could mask the CcnA binding regions. This final result consolidates the observation that CcnA may be indeed associated *in vivo* with the 5’UTR of *ctrA* expressed by the promoter P2 and not by P1. We also analyzed all possible interaction candidates bound to the 5’-UTR of *ctrA* P1 and P2 (Table S1). This analysis revealed that several other non-characterized ncRNAs might interact with the P1 and P2 5’-UTRs of *ctrA*. Their specific role should be investigated in future studies.

### CcnA interacts *in vitro* with CtrA and GcrA mRNAs

MAPS revealed a putative interaction between CcnA and P2-5’-UTR of *ctrA* and interestingly, among master regulators of cell cycle, the *gcrA* mRNA (Table S1). To better characterize/validate these interactions, we performed *in vitro* probing experiments. Results showed two regions of protection by CcnA for CtrA 5’UTR from the promoter P2 (Figure 4A). Concerning the *gcrA* mRNA 5’UTR, we used data derived from 5’ RACE experiments at the genome scale (Zhou et al., 2015). Results obtained with *in vitro* probing experiments for *gcrA* 5’-UTR instead showed only one region of protection by CcnA (Figure 4B). A common feature of both *gcrA* and *ctrA* P2 protections by CcnA was the sequence 5’-GGGG-3’ (Figure 4A, B) that corresponds to the region of CcnA belonging to a loop (Loop A) (Figure 4C). EMSA experiments using P2-*ctrA* and *gcrA* 5’-UTRs confirmed the interaction with CcnA (WT). The binding is diminished with a CcnA mutated in the Loop A (CcnA_GGGG_) (Figure 5A, B, C, D, E, F). We also performed EMSA on mutated P2-*ctrA* (P2-*ctrA_CCCC_*) and *grcA* (*gcrA_CCCC_*) and there was also a decrease of the CcnA binding (Figure S2B, C). However, CcnA_GGGG_ was not able to compensate the mutations on P2-*ctrA* or *grcA* 5’-UTRs (data not shown). Considering the putative importance of the Loop A for the interaction between CcnA, *ctrA* and *gcrA* mRNAs we searched for the presence of the GGGG motif in the 5’-UTRs of MAPS targets (Table S1) in comparison with a dataset of UTRs randomly selected in the genome of *C. crescentus*. Results showed that 35% of CcnA-bound MAPS positive candidate targets possessed GGGG (p-value = 0.02). As a stretch of CCCC, present in the Loop A region of CcnA, is protecting a putatively conserved GGGG motif in P2 *ctrA* 5’UTR and *gcrA* 5’UTR, we constructed mutant of CcnA of the Loop A by introducing mutations in the active loop “CCCC to GGGG” (CcnA_GGGG_). This mutated version of CcnA was then tested *in vivo* using the same pSRK expression system as previously. The mutation CcnA_GGGG_ in Loop A reduced the growth defect phenotype of CcnA overexpression (Figure 6A, B, C). Flow cytometry analysis of the Loop A mutant revealed a dominant negative phenotype similar to the antisense expression with accumulation of chromosomes (n≥3) (Figure 6D, E). These results suggest that the growth defect phenotype observed when inducing the WT version of CcnA could be mainly due to the interaction of the Loop A of CcnA to the mRNAs of *gcrA* and *ctrA*. As the interaction between CcnA and GcrA was confirmed *in vitro* we asked whether this binding was suggesting a possible regulation of CcnA on the GcrA protein levels. Therefore, we used the overexpression of CcnA and measured the level of GcrA using Western blot and anti-GcrA antibodies. The analysis of GcrA in a *ccnA* overexpression strain revealed a decrease of GcrA steady state protein levels in comparison with a WT strain carrying the empty vector (Figure S14B), suggesting the presence of either a CtrA-mediated inhibition of *gcrA* transcription and, in addition, a direct effect on GcrA expression by CcnA binding to its mRNA. However, besides showing a direct interaction between CcnA and the 5’UTR of *gcrA*, we are not able to disentangle the effect of CtrA regulation on GcrA activity from a potential direct regulation of the *gcrA* mRNA by CcnA.

**Figure 4.**
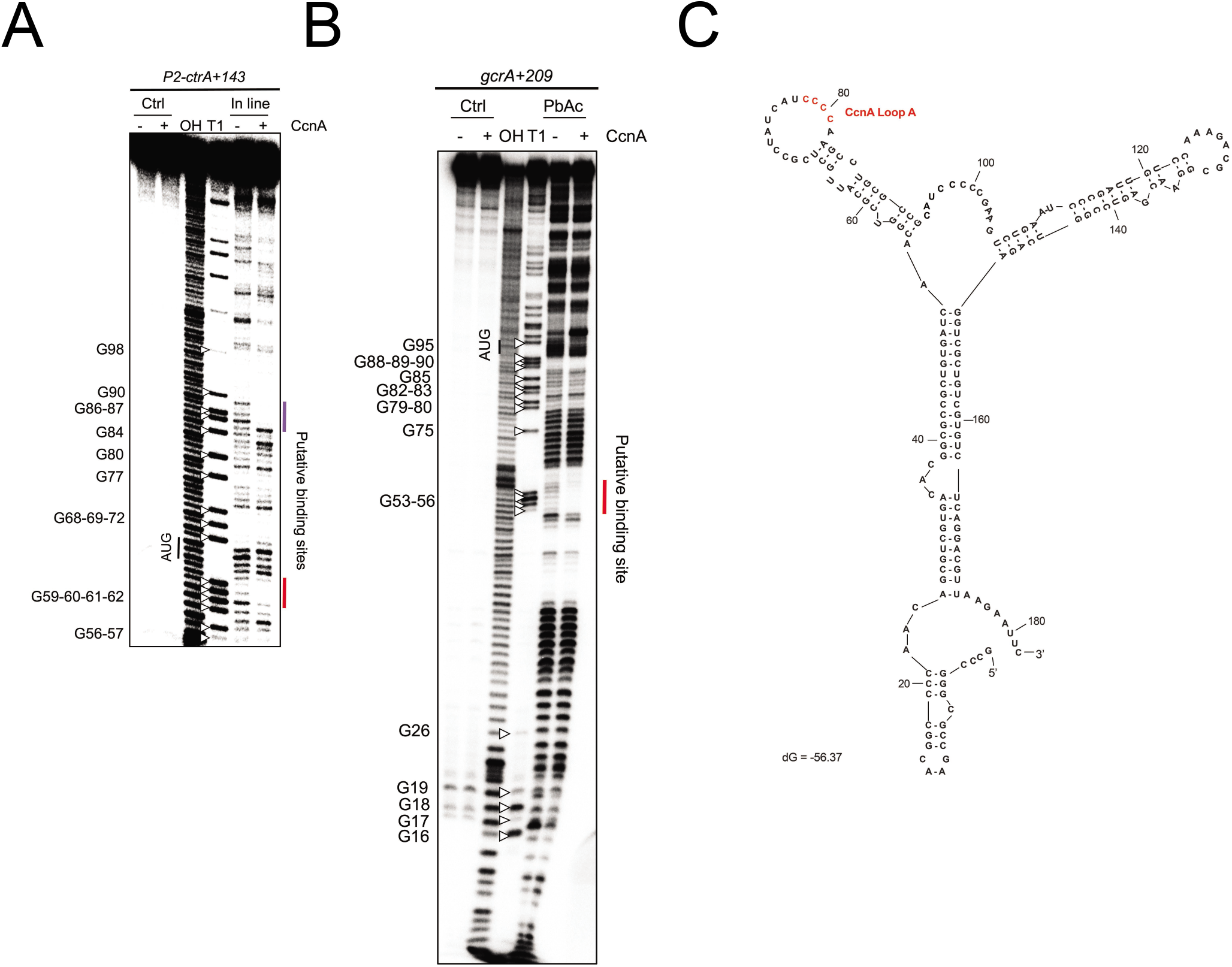
CcnA interacts directly *in vitro* with the 5’UTR of P2-*ctrA* mRNA and 5’UTR of *gcrA* mRNA. **A.** In-line (MgCl_2_) probing of 5’end-radiolabeled P2-*ctrA*+143 incubated in presence (+) or absence (-) of CcnA ncRNA. OH, alkaline ladder; T1, RNase T1 ladder. The numbers to the left indicate sequence positions with respect to the +1 of the transcript. **B.** Lead acetate probing of 5’end-radiolabeled *gcrA*+209 incubated in presence (+) or absence (-) of CcnA ncRNA. OH, alkaline ladder; T1, RNase T1 ladder. The numbers to the left indicate sequence positions with respect to the +1 of the transcript. **C.** Secondary structure of CcnA RNA predicted using mFold algorithm (Zuker, 2003). The predicted free energy of the thermodynamic ensemble is −56.37 kcal/mol. CcnA Loop A is shown in red and is composed of a stretch of « CCCC »

**Figure 5.**
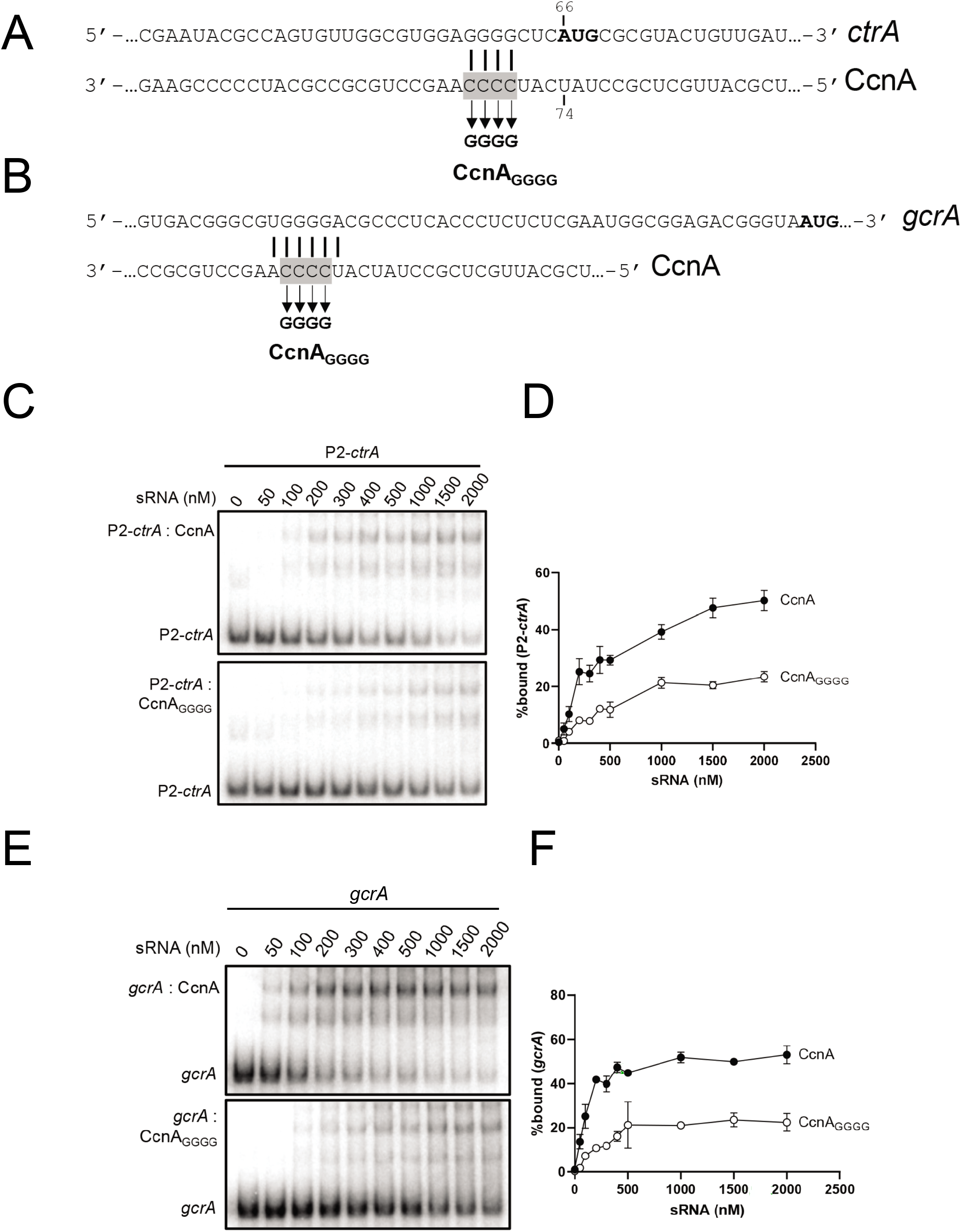
CcnA mutation of Loop A decreases the interaction with the 5’UTR of P2-*ctrA* mRNA and 5’UTR of *gcrA* mRNA. **A.** Mutation of the *ctrA* mRNA (*ctrA_GGGG_*) and ncRNA CcnA (CcnA_GGGG_) binding site (shown in red in Figure 4A). Solid lines indicate CcnA binding sites on *ctrA*. Boxed gray text corresponds to the nucleotides mutated. The translation start codon is shown in bold. **B.** Mutation of the *gcrA* mRNA (*gcrA_GGGG_*) and ncRNA CcnA (CcnA_GGGG_) binding site (shown in red in Figure 4B). Solid lines indicate CcnA binding sites on *gcrA*. Boxed gray text corresponds to the nucleotides mutated. **C.** 5nM of P2-*ctrA* (+143nt from the P2-*ctrA* promoter) RNA fragment was incubated with increasing concentration of CcnA (top) or CcnA_GGGG_ (bottom) ncRNA. % bound RNA with ncRNA CcnA (black) or CcnA_GGGG_ (white) is showed (**D**). Data represent the mean of two independent experiments ± SD. **E.** 5nM of *gcrA* RNA fragment was incubated with increasing concentration of CcnA (top) or CcnA_GGGG_ (bottom). % bound RNA with ncRNA CcnA (black) or CcnA_GGGG_ (white) is showed (**F**). Data represent the mean of two independent experiments ± SD.

**Figure 6.**
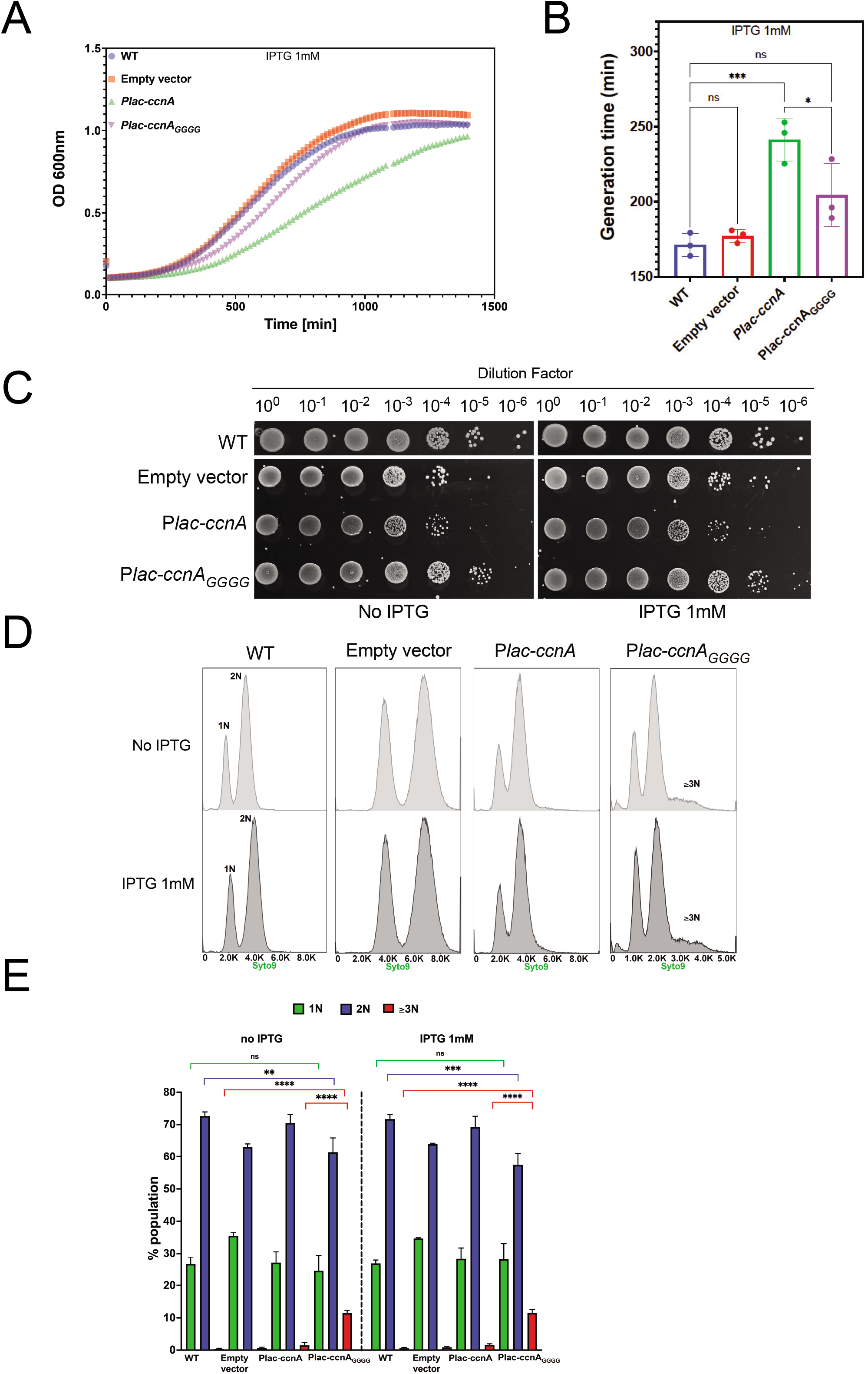
CcnA Loop A mutant shows attenuated cell cycle defects. **A. Growth curves following the expression of CcnA mutated in its Loop A (P*lac-ccnA_GGGG_*).** WT cells and WT cells carrying either an empty pSRK (Empty vector) or a pSRK with *ccnA* under the control of an inducible P*lac* promoter (P*lac-ccnA*) or *Plac-ccnA_GGGG_* mutated in its loop A were grown overnight in PYE without IPTG. 200μL of cells back-diluted from stationary phase cultures to an OD_600nm_= 0.02 were grown on 96 wells in PYE supplemented with 1mM IPTG. Cell growth was monitored overnight with a Spark-TM at 30°C and a shaking (orbital) amplitude of 6 mm and a shaking (orbital) frequency of 96 rpm. Results are shown as mean N=3 biological with 3 technical replicates. Raw data are provided in Table S7. Note that all growth curves data of figure 2, 6 and S9 were acquired in the same days of their respective biological replicates and compared to each other. **B. Determination of the doubling time of cells expressing P*lac-ccnA_GGGG_*.** Doubling times of cells from (6A) were calculated by using the exponential growth equation (Nonlinear regression) (Prism GraphPad 9.1.2). Statistical analysis was performed using ANOVA with a Brown-Forsythe and Welch test with a Dunnett’s multiple comparisons test. Ns = difference not significant, *: p.val =0,0396 ***: p.val = 0,0009. Raw Data are provided in Table S7. **C.** WT cells, WT cells carrying an empty vector, *Plac-ccnA* or *Plac-ccnA_GGGG_* were grown overnight in PYE at 30°C and diluted to an OD_600nm_= 0.6. Samples were then serially diluted (10^0^–10^−6^) and 4.5 μL of each dilution were spotted on a PYE-Agar plate with or without IPTG 1mM and incubated at 30°C. WT cells without plasmid were used as negative control. **D.** Flow cytometry profiles after SYTO 9 staining showing DNA content of WT cells, WT cells carrying either an empty pSRK (Empty vector) *ccnA* under the control of a *Plac* promoter (P*lac-ccnA*) or *ccnA* Loop A variant (*Plac-ccnA_GGGG_*) grown in PYE until OD_600nm_ = 0.3. Then induction of P*lac-ccnA* or P*lac-ccnA_GGGG_* was made by addition of IPTG 1mM 30min. Cells without induction were grown for an additional 30min as a control of growth phase. A total number of 300 000 particles were analysed by flow cytometry. **E.** Proportions of cells harboring 1N, 2N and ≥3N DNA in the population were analysed by gating the histograms in E). Data are representative of 3-5 biological replicates. Statistical analyses were carried out using ANOVA Tukey test. ns: difference not significant, **: p.val <0.01, ***: p.val <0.001, ****: p.val <0.0001.

Taken together those results suggest that the region corresponding to Loop A plays a significant role in the CcnA activity, confirming both the *in vitro* and the MAPS results shown previously. However other regions can definitely play important roles in the activity of CcnA that will require further analysis. Moreover, CcnA seems to have a second important target in the cell, GcrA, for which the ncRNA plays a negative role.

### CcnA affects the CtrA and GcrA regulons

RNAseq was used to compare the strains overexpressing *ccnA* to the strains expressing *ccnA* antisense (*ccnA-as*) in biological triplicates in order to reveal RNAs affected by CcnA with the hypothesis that it may show links with CtrA and GcrA regulons.

Differentially expressed genes identified when comparing the sense and antisense expressing strains were considered for the analysis (Figure S12 and Table S2). These results were also integrated with additional information such as (i) the presence of full or half CtrA binding boxes as identified by a Position Weight Matrix scan of the *C. crescentus* genome (Brilli et al., 2010a), (ii) the abundance of reads from a ChIP-Seq experiment aimed at characterizing GcrA occupancy (Haakonsen et al., 2015a), (iii) the genes whose expression levels change significantly in a Δ*ccrM* strain (Gonzalez and Collier, 2013), (iv) the essential genes as revealed with Tn-seq (Christen et al., 2011a), (v) genes with cell cycle-dependent expression (Fang et al., 2013a). The analysis revealed 215 genes differentially expressed in the two strains (CcnA vs CcnA-as). The CtrA regulon is composed of genes activated and repressed by the phosphorylated form of CtrA, which recognizes a full palindromic or half site (Zhou et al., 2015). Among the 215 genes, we found a statistically significant enrichment of CtrA binding sites, both half and full (Brilli et al., 2010a). To calculate significance of enrichments, we used a one sided binomial exact test (binom.test in R) and got a p-value=0.0065 for the full site, and a p-value=0.0001 for the half site. This finding suggests that upon changes of CcnA levels, the most affected regulon is CtrA’s.

We also looked for differentially expressed genes that could be part of the GcrA regulon. Many genes identified contained a GcrA binding region, suggesting that the GcrA regulon is differentially modulated in presence (*ccnA*) or absence (*ccnA-as*) of CcnA. Most of the genes of figure S12 are cell cycle-regulated as expected considering that both GcrA and CtrA are controlling those genes (Figure S12). Therefore, RNA-seq allows getting a full picture of the effects of CcnA activity perturbations which affect a significant fraction of the transcriptome involved in cell cycle regulation.

We also tested if genes affected by overexpression and inactivation of CcnA were also directly interacting with CcnA, as revealed by MAPS analysis (Figure S12, Table S1). Results showed that several genes that change expression levels upon mutations of CcnA are in fact putative direct targets of the ncRNA, for example, the mRNA encoding the transcriptional regulator MraZ, involved in the cell division processes, the GGDEF diguanylate cyclase DgcB, the RNA polymerase sigma factor RpoH or the polar organelle development protein PodJ.

In conclusion, the RNAseq results consolidate the potential effect of CcnA on CtrA and GcrA regulons as direct regulator of CtrA and GcrA protein levels but also showing a CcnA link with genes controlled by those master regulators as revealed by MAPS. Moreover, the overexpression of *ccnA* antisense shows opposite effects than the overexpression of wild type *ccnA*.

### Overexpression of CcnA complements cell cycle defects

As CcrM-dependent adenosine methylation sites (G<u>A</u>nTC) are connected to *ctrA* transcription by its own P1 promoter, we asked whether the expression of CcnA (or its antisense) was rescuing the Δ*ccrM* mutant cell cycle severe phenotype (Murray et al., 2013), considering that CcrM methylation is required to recruit GcrA at the P1 promoter region and therefore activate *ctrA* transcription (Mohapatra et al., 2020) (Figure S13). We attempted to introduce the plasmid containing *ccnA* and *ccnA-as* in the *ΔccrM* mutant and analyzed the different phenotypes. First, we were unable to introduce the plasmid carrying *ccnA-as* into Δ*ccrM*, suggesting an incompatibility between the two genetic constructs, while the electroporation frequency for wild type was as expected. This can be explained considering that both CcrM and CcnA are important to properly express CtrA, therefore, removing both mechanisms may be lethal. On the contrary, the expression of CcnA in Δ*ccrM* was viable and indeed able to suppress cell cycle defects of the mutant (Figure 7A). Notably, the severe morphological defects of Δ*ccrM* were rescued (Figure 7A, B, C), as well as the motility defects (Figure 7D). We also noticed that Δ*ccrM* cells rescued by CcnA were not curved (Figure S14A). This suggests that Δ*ccrM* still retains some of the features that are independent from CtrA, as the cell curvature depends on the gene *creS* encoding for the crescentin responsible for the methylation-dependent cell curvature of *C. crescentus* (whose expression depends on GcrA) (Mohapatra et al., 2020).

**Figure 7.**
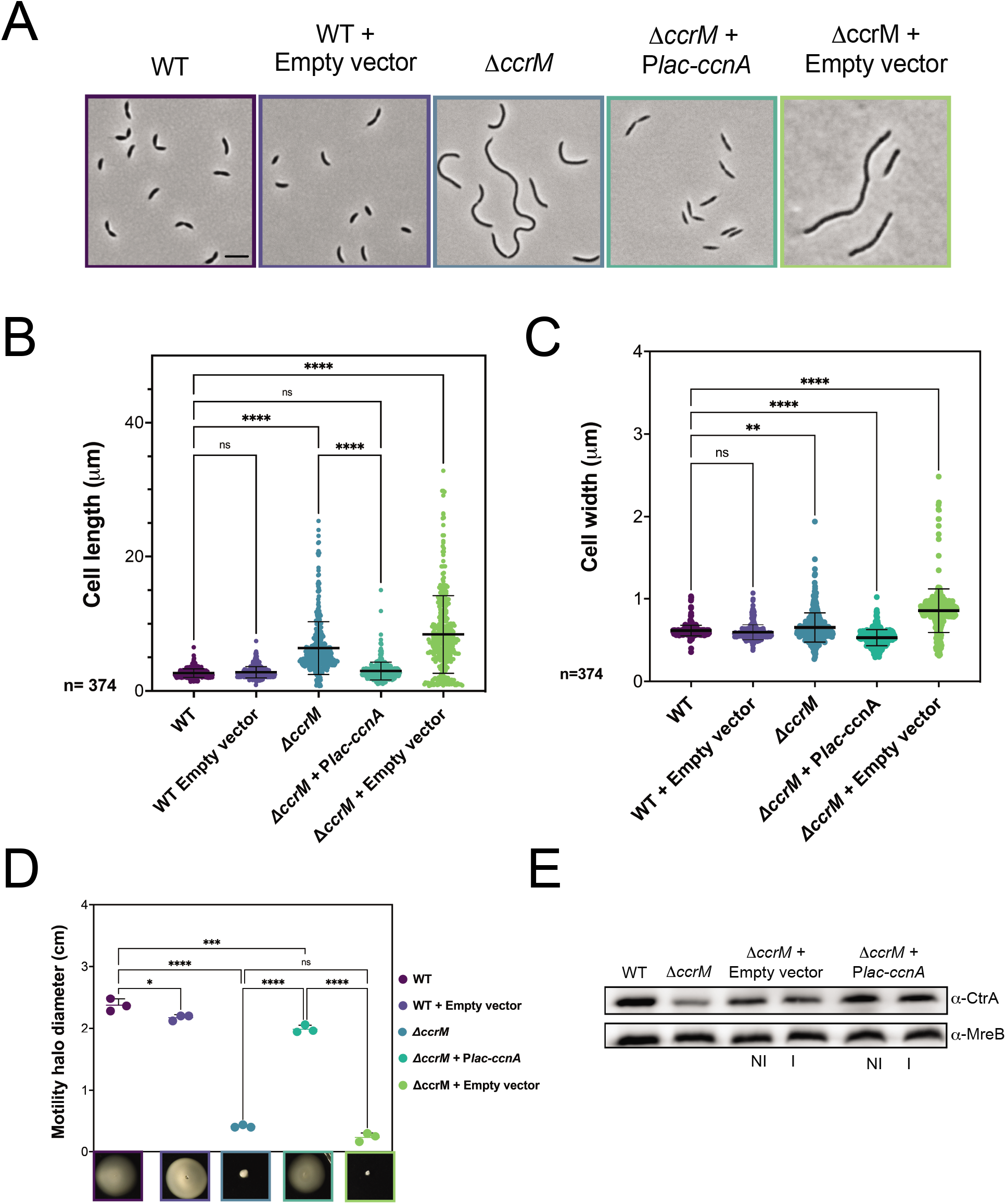
CcnA rescues the CcrM loss of function phenotype. **A.** Phase contrast images of WT cells, WT cells carrying an empty pSRK (empty vector), Δ*ccrM* cells, Δ*ccrM* carrying a plasmid with *ccnA* under the control of a *Plac* promoter (Δ*ccrM*+ P*lac-ccnA*) or Δ*ccrM* cells carrying an empty vector (Δ*ccrM+* empty vector) grown in PYE at 30°C until OD_600nm_= 0.6. Scale bar = 2μm. **B. C.** Cells from (7A) were analyzed using MicrobeJ (Ducret et al., 2016a) and 374 cells were analyzed to assess cell length and cell width. Statistical significance was determined using ANOVA with Šídák’s multiple comparisons test. ns: difference not significant **: p.val =0.0050 ****: p.val <0.0001. Raw Data are provided in Table S8. **D. Swarming assay on 0.25% soft-agar plates.** 1μL of each culture from cultures of figure 7A was deposited into the soft-agar and incubated at 30°C for 5 to 6 days. N=3. The diameter in cm of each mobility halo was measured with Fiji and reported in Table S4. Statistical significance was determined using ANOVA with Šídák’s multiple comparisons test. ns: difference not significant *: p.val=0,0406, ***: p.val= 0,0003, ****: p.val <0.0001. **E.** WT cells, Δ*ccrM* cells, Δ*ccrM* cells carrying an empty vector or P*lac-ccnA* were grown in PYE at 30°C until OD_600nm_ = 0.6. Then, induction of P*lac-ccnA* was made by addition of IPTG 1mM 30min. As a control of induction Δ*ccrM* cells carrying an empty vector were also incubated 30min in presence of IPTG 1mM. Proteins were extracted and separated on a SDS-PAGE gel for Western blotting. CtrA and MreB (loading control) proteins were revealed using specific polyclonal antibodies on nitrocellulose membranes.

We asked whether CcnA was indeed able to increase CtrA steady state levels in the Δ*ccrM* strain (Figure 7E). As most of the GcrA-CcrM dependent promoters, *ctrA* P1 is sigma-70 dependent, thus able to provide a basal level of transcription even in absence of methylation. Results clearly showed that CcnA can increase CtrA steady state levels in the Δ*ccrM* mutant closer to the wild type levels. Presumably, the lower level of CtrA depends on the amount of mRNA corresponding to P2 that may be lower in the Δ*ccrM* background. Moreover, the mechanism by which CcnA increases CtrA protein levels is independent from CcrM, possibly acting on the P2 promoter.

To provide a more complete characterization of CcnA role, we combined CcnA ectopic expression (sense or antisense) with Δ*pleC*, a mutant impaired in the negative control of DivK phosphorylation level. By considering that (i) DivK~P inhibits CtrA stability and activity, and (ii) that PleC is DivK’s phosphatase, CtrA levels in the Δ*pleC* mutant are low (Figure 8A). Therefore, overexpression of CcnA might compensate the defects in this mutant, restoring a phenotype resembling the wild type.

**Figure 8.**
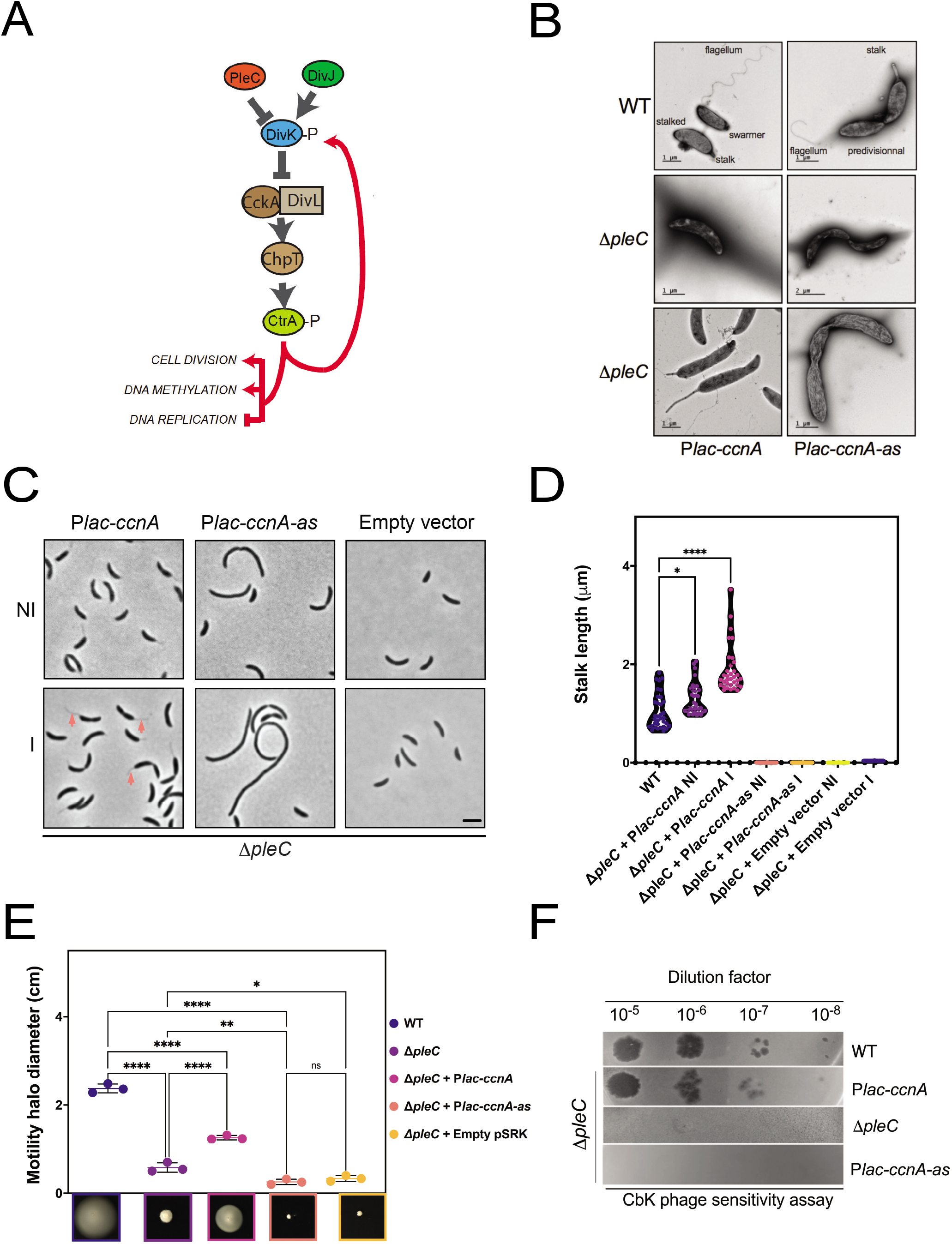
CcnA rescues the pleiotropic phenotypes of Δ*pleC*. **A.** Schematic representation of CtrA-DivK negative feedback loop in *C. crescentus*. DivK phosphorylation level is controlled by its kinase DivJ and its phosphatase PleC. At the swarmer cell pole, DivK must be dephosphorylated in order to enable the phosphorelay CckA-ChpT-CtrA. At the stalk pole, the presence of DivJ/absence of PleC keeps DivK fully phosphorylated leading to a block of CckA scaffold DivL. The absence of PleC causes a decrease of CtrA (Coppine et al., 2020), both at the phosphorylation and protein levels as CtrA phosphorylated (CtrA-P) controls its own transcription from the promoter P2. **B.** Electron microscopy images of WT cells, Δ*pleC* cells *and ΔpleC* cells carrying either P*lac-ccnA* or P*lac-ccnA-as* cells grown in PYE without IPTG at 30°C until OD_600nm_= 0.6. **C.** Phase contrast images of Δ*pleC* cells carrying a plasmid with *ccnA* or its antisense *ccnA-as* under the control of a *Plac* promoter (Δ*pleC*+ P*lac-ccnA* or Δ*pleC*+ P*lac-ccnA-as*) or Δ*pleC* cells carrying an empty vector (Δ*pleC+* empty vector) grown in PYE at 30°C until OD_600nm_= 0.6. Induction of *ccnA* or *ccnA-as* was made when cells reached 0.6 by the addition of IPTG 1mM for 30min. Scale bar = 2μm. **D.** Violin plots of stalks length per cell for each strain tested in Figure 8C plus a WT *C. crescentus* as a control for normal stalk length. Stalk length was measured by using BacStalk software (Hartmann et al., 2020). Statistical significance was determined using ANOVA with Brown-Forsythe and Welch’s tests with a Dunnett’s T3 multiple comparisons test. *: p.val =0.0117; ****: p.val<0.0001. Raw data are in table S5. **E. Swarming assay on 0.25% soft-agar plates.** 1μL of each culture from cultures of figure 8C was deposited into the PYE soft-agar and incubated at 30°C for 5 to 6 days. N=3. The diameter in cm of each mobility halo was measured with Fiji and reported in Table S4. Statistical significance was determined using ANOVA with Šídák’s multiple comparisons test. ns: difference not significant, *: p.val=0.0242, **: p.val= 0.0039, ****: p.val <0.0001 **E. CbK phage sensitivity assay**. A bacterial layer of cultures from WT, Δ*pleC* + *ccnA*, Δ*pleC*, or Δ*pleC + ccnA-as* was deposited into a PYE-Agar plate and incubated at 30°C. CbK phages were serially diluted (10^0^ – 10^−8^) and 4.5 μL of each phage dilution were spotted on top of the cultures and incubated at 30°C to visualize cells lysis. WT and Δ*pleC* cells were used as a control of the presence or absence of lysis, respectively.

We introduced *ccnA* or *ccnA-as* in Δ*pleC* mutant and observed the morphology, motility in soft agar plates, sensitivity to the CbK phage and stalk length. In agreement with our reasoning, the ectopic expression of CcnA was able to rescue Δ*pleC* defects, restoring stalks and motility while the expression of the CcnA antisense caused a very severe phenotype (Figure 8B, C, D, E). Electron microscopy was used to characterize more in details the phenotypes (Figure 8B). Results showed that upon CcnA expression (Figure 8C), stalks were longer in the Δ*pleC* background cells compared to WT cells (Figure 8D) and motility was also partially restored (Figure 8E). On the contrary, the expression of the antisense induced a severe growth (data not shown) and morphological phenotype with absence of polar structures in the majority of cells (Figure 8C, D).

We asked whether this suppression was just obtained by increasing the level of CtrA or if it was also able to affect the phosphorylation, and therefore the activity of CtrA. We measured CtrA~P by Phos-Tag technique (Figure S15). This analysis revealed that the CcnA expression was indeed able to increase protein levels of CtrA and slightly CtrA~P in Δ*pleC*.

Finally, we measured the sensitivity of *C. crescentus* to the phage CbK, which is adsorbed by the flagellum and enters the cells by attachment to the pili structures (Figure 8F). As the main subunit PilA of the pilus is completely under the control of CtrA, a Δ*pleC* mutant has an unfunctional flagellum and no pili, making this strain resistant to CbK infection (Panis et al., 2012; Sommer and Newton, 1988). Results showed that the expression of CcnA was able to completely restore the sensitivity of *C. crescentus* to CbK to WT levels, suggesting a *de novo* synthesis of the pili. The expression of CcnA-as did not change the resistance to the phage infection of the Δ*pleC* mutant, as shown by phage-induced lysis (Figure 8F).

### Conservation of CcnA in the class *Alphaproteobacteria*

Considering the key role of CcnA in *C. crescentus* coordinating CtrA and GcrA, two of the principal master regulators of cell cycle, we asked whether its function was conserved in bacteria that share the regulatory mechanisms by those master regulators. We considered a well-known bacterial model, *Sinorhizobium meliloti*, a symbiotic nitrogen-fixing organism. *S. meliloti* shares with *C. crescentus* most of the regulatory circuit driving cell cycle, including CtrA (Pini et al., 2013, 2015). Therefore, we took advantage of the expression system we used for *C. crescentus*, which is compatible with expression in *S. meliloti* (Khan et al., 2008). Expressing *C. crescentus* CcnA in *S. meliloti* slowed growth and caused an abnormal cellular morphology (Figure S16A) in comparison with the same vector expressing the empty plasmid. We therefore asked whether this alteration in cell morphology was due to a change in CtrA steady state levels (Figure S16B). Indeed, the overexpression of *ccnA* in *S. meliloti* cells showed an increase of CtrA proteins levels in comparison with the strain containing the empty vector, suggesting a similar mechanism than *C. crescentus*. Results showed that CcnA of *C. crescentus* is able to induce a cell cycle defect, that is branched cells and a clear cell division retard, similar to that observed in a delta-*divJ* mutant (Pini et al., 2013) and presumably linked to an increased level of CtrA.

The activity of *C. crescentus* CcnA in these two alphaproteobacterial species suggested that a putative homologous gene should be present in *S. meliloti*. We therefore scanned the genomes of the alphaproteobacterial species using GLASSgo (Lott et al., 2018a) aiming to find CcnA homologs. We found a conservation of CcnA in several closely related species (Figure S16C). As expected CcnA has closer homologs in the *Caulobacterales*, but it can also be found in the other families except for the *Rickettsiales*. Considering that *Rickettsiae* have experienced a massive reduction of the genome, it is reasonable to speculate that CcnA may be a conserved factor that has coevolved with CtrA, participating in the ancestors to its regulation of transcription. Taken together these results prompted us to compare 5’UTRs of *ctrA* in these two organisms in order to find shared motifs potentially complementary to CcnA sequence and in conclusion involved in *ctrA* translation. By using an *in-silico* analysis made with the Clustal Omega software (Madeira F et al., 2019), we found that the stretch of GGGG putatively interacting with CCCC of CcnA within its Loop A is conserved in the *ctrA* 5’UTR of *S. meliloti* separated from the start codon by 6 nucleotides instead of 3 for *C. crescentus* CcnA (Figure S16D). This may explain why CcnA from *C. crescentus* is able to increase CtrA protein level in this species, even if a “CcnA-like” homolog was not clearly detected in *S. meliloti*.

## Discussion

The origin of replication of *C. crescentus* is necessary for replication of the chromosome and therefore represents one of the most important regions of the genome. CtrA binding sites at the origin of replication play an inhibitory role on the replication of DNA as they allow CtrA~P to compete out the binding of DnaA (Frandi and Collier, 2019). Transcriptomic data indicated that some parts were nonetheless transcribed; in particular, a short gene was found transcribed (CCNA_R0094), corresponding to an essential genome region highlighted by the analysis of TnSeq data (Christen et al., 2011a; Schrader et al., 2014; Zhou et al., 2015). This gene is surrounded by CtrA boxes at −23 bp from the TSS and at the very end of the gene (Brilli et al., 2010a). In the process of understanding the role of this non-coding RNA, belonging to the origin of replication, named here CcnA, we found that CcnA is a regulator of cell cycle, specifically linked to two master regulators, CtrA and GcrA. To the best of our knowledge this is one of the first demonstrations of a ncRNA playing a stress-independent role in the cell cycle regulation of a bacterium. Examples of regulatory ncRNAs controlling key cellular functions can be found elsewhere in addition to the nowadays classical RyhB pathways controlling iron utilization in *E. coli*, such as the Qrr ncRNAs in *Vibrio* species, that participate in quorum sensing, or NfiS, a positive regulator of the Nitrogenase in *Pseudomonas stutzeri* A1501 (Zhan et al., 2016), which is folded into a compact structure that acts on the mRNA of *nifK*, encoding the β-subunit of the MoFe protein of the nitrogenase enzymatic complex, enhancing its translation.

Using qRT-PCR, we clearly showed that CcnA starts accumulating in the second half of the S-phase, coincidentally with the accumulation of CtrA, presumably as an effect of *ctrA* transcription from its promoter P1. Using several approaches, we hypothesized that expression of *ccnA* depends on cell cycle, presumably by CtrA. We also found that once CcnA starts to accumulate, it binds the mRNA of *ctrA* by base pairing using at least one region belonging to a loop predicted to exist in its structure (Figure 4C). *In vitro* probing experiments on *ctrA* and *gcrA* 5’UTRs showed that a stretch of CCCC is particularly important for CcnA to interact with its target mRNAs, possibly stabilizing the interactions. We hypothesize that this binding of CcnA on the *ctrA* UTR frees the RBS enabling translation at higher rates and therefore causes an increase in the protein levels. We predicted the structure of the UTR starting from the TSS of promoter P2 of the gene *ctrA* and it appears evident that the mRNA of CtrA has its putative Shine-Dalgarno (SD) of the RBS at −6 from ATG sequestered in a stem (Figure 9A). Although classically, ncRNAs pairing at the SD induce translational block, which is in disagreement with our observations, probing revealed another region of the *ctrA* mRNA that is impacted in presence of CcnA (Figure 4A), which is more compatible with a positive regulation of *ctrA* translation by CcnA. It has already been shown that a pairing of a ncRNA at the beginning of the coding sequence can have an activating role (Jagodnik et al., 2017). Hence, we can imagine that both binding are important and both responsible for the role of CcnA on *ctrA*. We attempted to construct a CcnA mutant corresponding to this interaction. Unfortunately, the introduction of this mutation in CcnA makes the RNA unstable. (data not shown). Future studies on the structure of the *ctrA* UTR and CcnA may help elucidating this unorthodox positive mechanism of activation.

**Figure 9.**
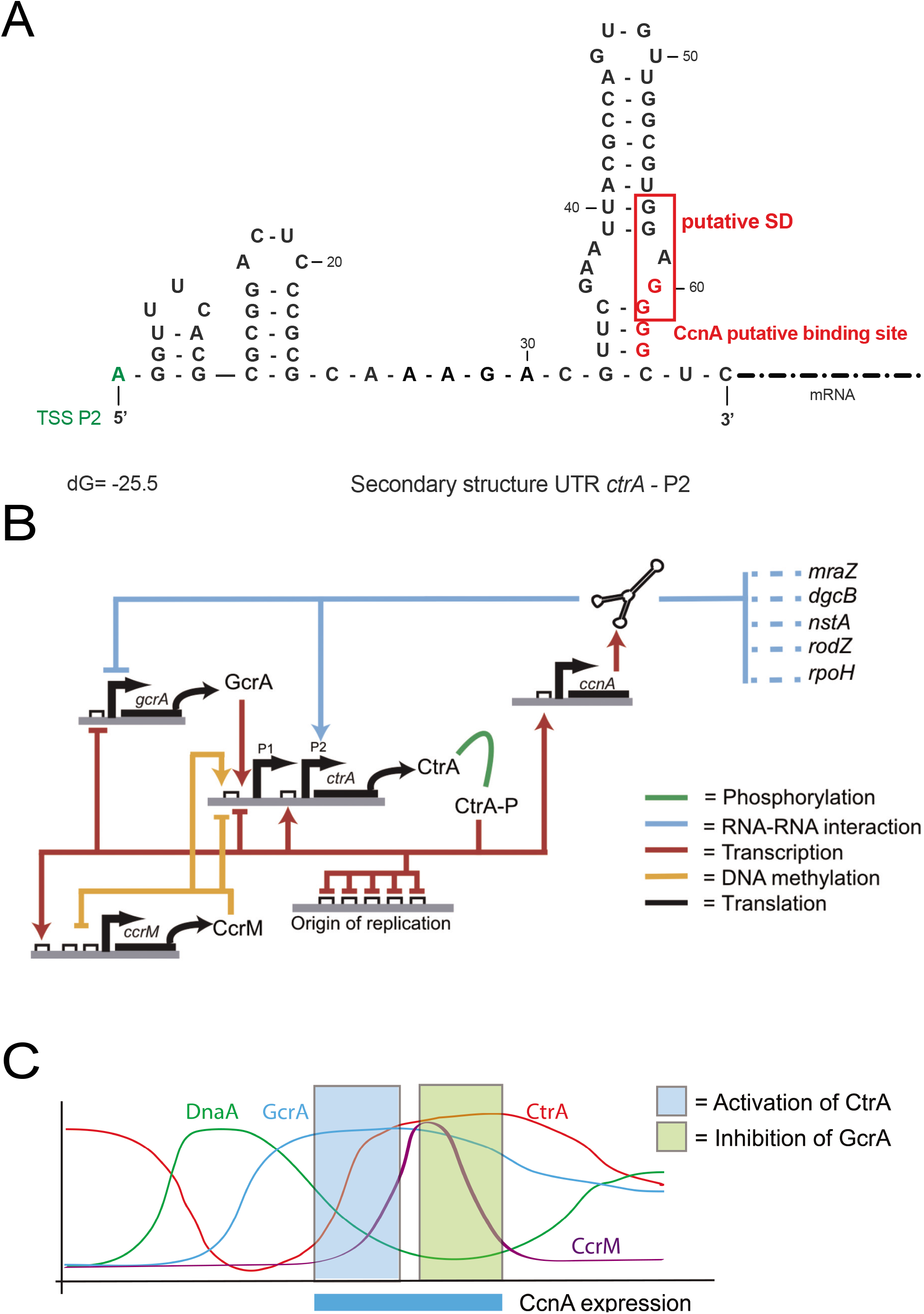
Integration of CcnA in the cell cycle regulation model in *C. crescentus*. **A**. Secondary structure prediction of 5’UTR of *ctrA* starting from the TSS of *ctrA*-P2 using « The DINAmelt Web server » – Two-state melting (folding) with default parameters for RNA energy rules (Markham and Zuker, 2005). The predicted free energy of the thermodynamic ensemble is −25.5 kcal/mol. “GGAGG” *ctrA*-P2 RBS is framed in red and appears to be blocked in a stem loop. Binding site of CcnA Loop A is indicated in red. **B.** Throughout the cell cycle the cascade of transcriptional activation of the gene *ctrA* involving GcrA and CcrM activates *ctrA*-P1 expression leading to its first protein accumulation. After the translation and activation by phosphorylation of CtrA, CtrA~P will reach the origin of replication to inhibit DNA replication. Our work suggests that simultaneously CtrA~P is potentially responsible of *ccnA* transcription. CcnA in return will create a positive feedback loop on CtrA protein accumulation after its P2 expression. This suggests that CcnA may be a key element of the second strong CtrA accumulation during the cell cycle. CcnA may also be a « CtrA-activity » modulator as its other putative targets belongs to the CtrA regulon. **C.** Concomitantly, CcnA regulates negatively putatively the translation of *gcrA* mRNA leading to a decrease of GcrA and presumably a correct and precise shut ON or OFF of the two master regulators. CcnA cell cycle expression window correlated in space and time with the activation and inhibition of CtrA and GcrA respectively. CcnA is proposed to act negatively on *gcrA* mRNA translation avoiding a *de novo* transcription of *ctrA-P1* and at the same time positively on *ctrA*-P2 mRNA translation to regulate the second wave of CtrA activation necessary for the expression of genes involved in fundamental processes such as cell division, chemotaxis, DNA methylation and biogenesis of polar structures.

An intriguing question about CcnA is its functional relationship with the origin of replication. Why does CcnA belong to the origin of replication? It is fascinating to speculate that CcnA belongs to the *CORI* as it must be fired at low levels of CtrA~P, therefore “using” high affinity CtrA binding sites (Taylor et al., 2011). This allows the presence of CcnA when the second mRNA of *ctrA*, generated from the P2 promoter starts accumulating. CcnA may be potentially involved in the translation of P2 mRNA of CtrA and therefore may act as a cell-cycle timer through CtrA activation (Kaczmarczyk et al., 2020).

Indeed, CcnA plays a role in the regulation of the expression of CtrA as a putative activator of translation. In our model (Figure 9B, C), the regulatory circuit created by CtrA-CcnA and back to CtrA represents a positive feedback loop in which the regulatory layer controlled by CcnA acts on top of a second layer of transcriptional auto-activation of *ctrA* on its second strong promoter P2. In parallel, CtrA has a potential inhibitory activity on *gcrA*, creating a negative feedback loop in which GcrA activates CtrA, which in turn blocks GcrA. CcnA acts as well on this feedback reinforcing a reduction of translation by direct binding onto the 5’UTR of *gcrA*. Therefore, CcnA does not create new connections between master regulators of cell cycle but in fact acts on a preexisting circuit, presumably increasing the robustness of the system. This behavior by ncRNAs has been described before (Dutta and Srivastava, 2018; Mandin and Guillier, 2013; Nitzan et al., 2017). The role of ncRNAs is therefore to consolidate the robustness of transcriptional circuits by introducing a fast post-transcriptional control on the mRNAs produced by transcription factors. From this point of view CcnA may indeed act as key trigger for protein production linking transcription to translation. The importance of CcnA emerges when redundant mechanisms of CtrA control are not present, such as the absence of CcrM (primary activator of CtrA expression in the second half of S-phase). In all systems investigated so far, ncRNA-mediated regulations introduce a rich variety of dynamical responses, but these have mainly been studied in the case of negative regulation by the ncRNA on the target transcript. Among the peculiarities of ncRNA-mediated negative regulation in bacteria, previous studies have observed a threshold linear response of target abundance and the possibility of an ultrasensitive response in target abundance as a function of the relative transcription rate of the ncRNA and the target (Levine et al., 2007; Mitarai et al., 2009). Moreover, ncRNAs may act as a fine-tuning of the affinity for different targets, but their effects might also create, in complex networks, phenomena such as bistability and oscillations (Liu et al., 2011).

Is this CcnA dependent mechanism, controlled by CtrA itself, also conserved in organisms in which CtrA regulates the cell cycle? We studied *C. crescentus* CcnA in *S. meliloti*, where the role of CtrA has been investigated (Pini et al., 2013, 2015). In these two organisms CtrA is essential and controls key cell cycle functions such as cell division and DNA replication. Consistently with our hypothesis, the expression of *C. crescentus* CcnA causes the same molecular alterations as described here in *C. crescentus*. Although more molecular investigation of homologous ncRNA in other organisms must be explored, we can hypothesize that CcnA activity may be a conserved mechanism of the regulation of the cell cycle. This new system of complex regulatory circuits carried out by CcnA indeed expand the key role of ncRNAs in bacteria, opening a new activity that will need a thorough molecular investigation of mechanistic activity of this ncRNA. The CcnA structure and consequent activity may be a new class of ncRNAs whose role is still at its beginning of study. Interestingly, a prediction of target genes among several homologs (data not shown) have revealed that targets usually fall into the chemotaxis and motility classes of genes, suggesting a common function. This is not surprising considering that CtrA itself is considered, in *C. crescentus* and most of alphaproteobacterial species, as a regulator of motility (Brilli et al., 2010a; Greene et al., 2012).

In conclusion the regulatory mechanism centered on CcnA represents an archetype of regulatory architecture. CtrA autoregulates itself via its promoter P2 and inhibits the expression of GcrA via its binding site on the promoter region of *gcrA*. The same two connections are performed by CcnA that activates CtrA translation and inhibits GcrA expression. This module on top of a more classical transcriptional regulation presumably ensures a strong effect during cell cycle. Taking advantage of the simplicity of this bacterial system, more specific experiments must be performed in order to elucidate this network behavior.

## Materials and Methods

### Strains, growth conditions and molecular biology techniques

Strains used in this work are listed in 3. *C. crescentus* strains were routinely cultured in peptone-yeast extract (PYE) medium with appropriate amount of antibiotics (Solid: Kanamycin 25 μg/ml, Tetracycline 2 μg/ml, Spectinomycin 100 μg/ml) (Liquid: Kanamycin 5μg/ml, Tetracycline 1μg/ml, Spectinomycin 25 μg/ml) and 0.3% xylose or 0.2% glucose whenever necessary. *S. meliloti* strains were cultured in Tryptone-Yeast extract (TY) medium with appropriate antibiotics (Streptomycin 500 μg/ml, Kanamycin 200 μg/ml). *E. coli* was grown in Lysogeny Broth medium. The cultures were grown at 30**°**C or 37**°**C as required for different experiments. Synchronization of the *C. crescentus* cells was done using Percoll or Ludox as described before (Marks et al., 2010). *E. coli* strains were grown at 37**°**C in LB broth or solid medium with required amount of antibiotic supplements (Ampicillin 100 μg/ml, Kanamycin 50 μg/ml, Tetracycline 10 μg/ml) as necessary. *C. crescentus* cells were transformed with different plasmids by electroporation. Western blotting were performed as previously described (Pini et al., 2015) using antibodies against CtrA, DnaA, GcrA and MreB using 1:5000 dilutions. pSRK vectors were constructed as previously described using primers listed in table S3 amplified using the polymerase Q5 (NEB). Soft Agar plates were prepared with 0.25% agar; images were taken using an IC-Capture Camera at 75% of magnification. Phostag was performed as previously described (Pini et al., 2015). CbK phage sensitivity assay was also performed as previously described (Panis et al., 2012).

### MS2-affinity purification coupled with RNA sequencing

Strains containing MS2-CcnA and MS2 UTRs of *ctrA* P1 and P2 were induced by 1 mM IPTG for 30min harvested and used to perform MAPS as previously described (Lalaouna et al., 2017). Analysis was performed by the following protocol. Reads were mapped to the indexed *C. crescentus* NA1000 genome (NC_011916) with Bowtie2 (Langmead et al., 2018) by using the following command: “bowtie2 --qc-filter --threads 18 --no-mixed --mp 10 --no-discordant -x NA1000 --passthrough −1 R1_001.fastq.gz −2 R2_001.fastq.gz”, which only returns concordant alignments in the form of *innies* (mates face each other) with at least 10 of MAPQ score. As we wanted to align reads that also fall outside coding sequences, we first mapped on the genome and then we used genome regions defined as explained below to calculate the coverage of the regions. In doing so, we need to consider that the paired libraries were obtained by using a stranded protocol (Illumina). For this reason, we first split the genome alignments into two files, one containing all pairs assigned with the flag 99/147 and the other reads pairs with flag 83/163. Basically, in doing so, we are putting all pairs aligned on the genome with a certain orientation in one file and all those aligned in reverse orientation in another. Each file is used as input to BamCoverage (Ramirez et al., 2016) to calculate coverage in 1 nt bins of the genome. At this point we used several files containing genome region coordinates (described below) to calculate the coverage of the regions, by taking reads on the basis of the expected alignment orientation wrt the transcript. For instance, when considering CDS, we will proceed similarly to what is done in standard RNA-seq i.e. we will calculate the coverage of the CDS by summing all the genome coverage values that fall within the CDS in the expected orientation. In this way we were able to calculate the coverage of pre-defined regions that are not present in the annotation file (i.e. the gff) of the NA1000 genome. Once obtained the coverage for our regions, we analysed them independently, by calculating a log2 ratio of the normalized coverage in the MS purified sample and the control. We defined as candidate targets for CcnA all genes for which one of the regions have a log2 ratio of the coverage of at least 2 (4-fold increase) using the RPM transformed data. To avoid artifacts for small coverage values that are subject to high experimental fluctuations, we also ask that each region has a coverage larger than the lower 25% of the regions in the MS experiment.

Most tools developed to calculate sequencing coverage from RNA-seq data usually rely on a pre-existing genome annotation, and among all features encoded in that file, they often focus on “CDS” or “gene”. This can have problems, as for instance ncRNA do not have a CDS associated and therefore tools strictly using CDS coordinates will completely overlook ncRNAs. In the present context, we were interested in understanding if MAPS data might allow inference about more detailed questions concerning a small RNA target transcript. For instance, if we can get information on the specific region of the transcript that is bound by the sRNA under examination. Together with defining a list of potentially bound transcripts in the different MAPS experiments performed in this work, we also defined 5’- and a 3’ UTRs for each gene, and analysed the coverage of the three regions independently. The 5’ UTR of a gene was defined on the basis of the experimentally determined transcription starts sites from Zhou et al., 2015 if the gene was present in their data, else as the 100 nt region upstream of the gene. Similarly, to avoid considering short UTRs, if the UTR defined by Zhou et al., 2015 was less than 100nt, we define the 5’-UTR as the 100-nucleotide region preceding the start of the CDS or ncRNA. As there is no similar experimental data for 3’-UTRs, we arbitrarily defined these regions as the 250 nucleotides going from 50 nt within the CDS or ncRNA to 200 nt downstream.

### Microscopy analysis

Cells were observed on a 24×50 mm coverslip under a 0.15% agarose-PYE “pad” to immobilize the cells. Samples were observed thanks to an epifluorescent-inverted microscope Nikon Eclipse TiE E PFS (100 × oil objective NA 1.45 Phase Contrast). Cells morphologies and fluorescent images were analysed using ImageJ and MicrobeJ (Ducret et al., 2016a; Schneider et al., 2012). Stalk length was measured by using BacStalk software (Hartmann et al.). Electron microscopy (EM) was performed by placing 5 μL drops of the bacteria suspension for 3 minutes directly on glow discharged carbon coated grids (EMS). The grids were then washed with two drops of 2% aqueous uranyl acetate, and stained with a third drop for 2 min. Grids were dried on filter paper and the samples were analyzed using a Tecnai 200KV electron microscope (FEI), and digital acquisitions were made with a numeric camera (Oneview, Gatan).

### Quantitative Real-Time-PCR for Transcriptional Analyses

RNAs were prepared from cultures at OD_600_~ 0.6). The cells were harvested and frozen at −80°C. Total RNAs were isolated from the pellet using the Maxwell 16 LEV miRNA Tissue Kit (Promega) according to the manufacturer’s instructions and an extra TURBO DNase (Invitrogen) digestion step to eliminate the contaminating DNA. The RNA quality was assessed by Tape station system (Agilent). RNA was quantified at 260 nm (NanoDrop 1000; Thermo Fisher Scientific). For cDNA synthesis, 1 μg total RNA and 0.5 μg random primers (Promega) were used with the GoScript Reverse transcriptase (Promega) according to the manufacturer instruction. Quantitative real-time PCR (qRT-PCR) analyses were performed on a CFX96 Real-Time System (Bio-Rad). The reaction volume was 15 μL and the final concentration of each primer was 0.5 μM. The cycling parameters of the qRT-PCR were 98°C for 2 min, followed by 45 cycles of 98°C for 5 s, 60°C for 10 s. A final melting curve from 65°C to 95°C is added to determine the specificity of the amplification. To determine the amplification kinetics of each product, the fluorescence derived from the incorporation of EvaGreen into the double-stranded PCR products was measured at the end of each cycle using the SsoFast EvaGreen Supermix 2X Kit (Bio-Rad, France). The results were analyzed using Bio-Rad CFX Maestro software, version 1.1 (Bio-Rad, France). Based on beta-galactosidase data, fusing the *ccnA* promoter with the ORF of *lacZ*, we found that CcnA transcription is high with levels around 10^4^ Miller units. Therefore, the RNA16S gene (also highly expressed) was used as a reference for normalization. For each point a technical duplicate was performed. The amplification efficiencies for each primer pairs were comprised between 80 and 100%. All primer pairs used for qRT-PCR are reported in the table S3.

### Flow cytometry analysis

*C. crescentus* cells grown to exponential, stationary phase or synchronized were harvested and stored in 70% ethanol at −20°C until further use. DNA content of cells was analyzed with Flow cytometry by using the protocol as described in (Bergé et al., 2020) with slight modifications.

For synchronized cultures, a population of pure G1 cells (swarmer cells) was obtained by separation with density gradient with Percoll. Briefly, cells from an overnight culture were diluted to OD= 0.1 and grown to 0.5-0.6, then centrifuged 5min at 8000rpm at 4°C. The supernatant was removed and the pellet resuspended in 750uL of cold 1X M2-Salt and mixed with 700uL of cold Percoll and vortexed then centrifuged at 12000rpm at 4°C for 20min. The top band (predivisional and stalk cells) was removed and the bottom band (swarmer G1 cells) was collected and washed 3 times in cold M2-Salt. The cells were then resuspended in 2mL of pre-warmed PYE (30°C). 200uL of samples following the cell cycle were collected every 15min from t=0 to t=120min and stored in 70% ethanol and processed as described below.

Due to the small size of the bacteria *C. crescentus*, we used a threshold and a trigger with the SSC signal (side scatter). The density plots obtained (small-angle scattering FSC versus wide angle scattering SSC signals) were gated on the population of interest, filtered to remove multiple events and then analysed for the fluorescence intensity (FL1 525 / 30nm) of the DNA probe SYTO™ 9 Green Fluorescent Nucleic Acid Stain at a final concentration of 2uM in the buffer (10mM Tris-HCl 7.5; 1mM EDTA; 0.01% triton X100; 50mM Na-citrate). Proportion of cells harboring 1N, 2N and ≥3N DNA were analysed by gating the peaks of the Syto9 fluorescence histograms. Samples were run in the low-pressure mode (5-10K events/s). A total number of 300-500K particles were collected per sample. Data were acquired with a S3e cells sorter (Bio-Rad) using 488 and 561 nm lasers and were analysed and plotted using FlowJo v10.6. Data are representative of 3-5 biological replicates and statistical analyses were carried out with Prism.v8.2 using ANOVA test.

### Probing experiments

Templates for in vitro probing, containing a T7 promoter, were obtained by PCR amplification. Lead acetate degradation and In-line probing assays were performed as previously described (Lalaouna et al., 2015). In brief, 0.2 μM of *in vitro*-generated *gcrA+209* and P2-*ctrA+143, 5*’end-labeled were incubated with or without 1μM CcnA ncRNA. Radiolabeled RNA was incubated 5 min at 90°C with alkaline buffer or 5 min at 37°C with ribonuclease T1 (0.1 U; Ambion) to generate the Alkaline (OH) ladder and the T1 ladder, respectively. RNA was analyzed on an 8% acrylamide/7M urea gel.

### RNA sequencing

Cultures were harvested at 0.6 OD_600_ and frozen in liquid nitrogen as previously described (Pini et al., 2015). Total RNA was prepared using RNeasy Mini Kit (Qiagen). Ribosomal RNAs were removed using the Bacterial RiboZero (Illumina) and libraries for MiSeq (V3 cassette) were prepared using the Stranded True Seq RNAseq Kit (Illumina). For the analysis of figure S12, reads were mapped using the Galaxy platform (Afgan et al., 2016) by Bowtie2, reduced to 10 bp Reads Per Kilobase per Million mapped reads (RPKM) in a Bedgraph format by BamCoverage (Ramírez et al., 2014) and visualized by IGV (Robinson et al., 2011). For analysis shown in figure S12, read alignments were performed with bowtie2 (Langmead et al., 2012) and the following additional parameters: --no-discordant --no-mixed --no-unal –dovetail. The resulting sam file was first converted into a bam file with samtools (Li et al., 2009) and then used as input to HTSeq count (Anders et al., 2015). Abundance matrices for all annotated genes were assembled together after removal of tRNA and rRNA genes, and used for differential gene expression analysis by using the R package DESeq2 (Love et al., 2014). Selection of differentially expressed genes was based on the contrast among libraries from a strain expressing the sense ncRNA CcnA and the strain expressing the corresponding antisense ncRNA by applying the following thresholds: FDR<0.01. We did not filter at a log fold change threshold to let the DESeq2 algorithm exploits the estimation of dispersion to provide a full list of likely differentially expressed genes (DEGS). This resulted in 215 DEGS, ranging in absolute value from a log fold change of 0.48 to a maximum of 2.9. Most of the DEGS are upregulated (208, or 97% of the total). The differential gene expression analysis was integrated with a number of available information on the cell cycle of *C. crescentus*: essentiality data come from (Christen et al., 2011b); the list of genes significantly changing their expression level during the cell cycle is from (Fang et al., 2013b) and are based on a RNA-seq experiment comprising 5 time points during the cell cycle in triplicate; GcrA ChIP-Seq data come from (Haakonsen et al., 2015b). We downloaded the reads corresponding to the GcrA sample and mapped them on the NA1000 genome to obtain a coverage profile. This profile was used to get an average coverage for each gene by considering the window going from 200 nt upstream of the ATG of the gene to 50 nt within the coding sequences. Data concerning the dependence of genes from methylation come from (Gonzalez et al., 2014) and were identified on the basis of a microarray analysis of strains engineered through removal of the gene encoding the methyltransferase (*ccrM*). The presence of CtrA binding sites (full and half) is based on scanning the genome with the PWM obtained by (Brilli et al., 2010b) and a threshold of 70% of the maximum score, calculated as from (Brilli et al., 2010b). Moreover, a site was indicated as present for a gene if it was found in the 250 nt upstream of the ATG and a site was assigned to the closest gene. The heatmap figure was obtained with the R package Pheatmap and integrates the above annotations with gene expression data in the present work (Figure S12).

### Primer extension

Transcriptional +1 of CcnA ncRNA was determined by primer extension. Briefly, 10μg of total RNA was incubated with 2 pmol of radiolabelled primer (EM5194) and 0.5mM dNTPs for 5 min at 65°C, followed by 1 min on ice. Reverse transcription was initiated by adding ProtoScript II Buffer (1x), ProtoScript II (200units, NEB) and DTT (5mM). The reaction mixture was incubated at 42°C for 1 h. The enzyme was inactivated at 90°C for 10 min. The reaction was precipitated and then migrated on a denaturing 8% polyacrylamide gel. Gel was dried, exposed to phosphor screens and visualized using the Typhoon Trio (GE Healthcare) instrument.

### Electrophoretic mobility shift assays (EMSA)

EMSA were performed according to Morita *et al*. (1), with some modifications. 5’-end-radiolabeled *ctrAp2, ctrAp2_LoopA, gcrA* or *gcrA-LoopA* was heated for 1 min at 90°C and put on ice for 1 min. *ctrAp2* or *ctrAp2_LoopA* RNA was diluted at 5 nM in binding buffer (50 mM Tris-HCl pH 8.0, 25 mM MgCl_2_, 20 mM KCl, 12.5 μg/mL yeast tRNA), *gcrA* or *gcrA-LoopA* RNA was diluted at 5 nM in binding buffer II (10 mM Tris-HCl pH 7.0, 10 mM MgCl_2_, 1000 mM KCl, 12.5 μg/mL yeast tRNA) and mixed with CcnA or CcnA-LoopA at different concentrations (0-2000 nM). Samples were incubated for 15 min at 37°C and reactions were stopped by addition of 1 μL of non-denaturing loading buffer (1X TBE, 50% glycerol, 0.1% bromophenol blue, 0.1% xylene cyanol). Samples were resolved on native polyacrylamide gels (5% acrylamide:bisacrylamide 29:1) in cold TBE 1X and migrated at 50 V, at 4°C. Gels were dried, exposed to phosphor screens and visualized using the Typhoon Trio (GE Healthcare) instrument. Image Studio Lite software (LICOR) was used for densitometry analysis.

## Acknowledgments

We thank members of the Biondi and Massé’s laboratory for critical comments on the manuscript. We thank the IMM Transcriptomic facility for the RNA preparation and the qRT-PCR experiment; we also thank Artemis Kosta and Hugo le Guenno from the IMM Microscopy platform for Electron Microscopy acquisition and analysis. We thank also Gaël Panis and Patrick Viollier for the phage CbK and also for providing the delta *cpdR, rcdA, popA* strains used in this work. We thank Regis Hallez and Romain Mercier for MreB and GFP antibodies, respectively, used in this study.

## Supplemental tables and figures

Table S1. MAPS and reverse MAPS targets.

Table S2. RNAseq of strains overexpressing *ccnA* and *ccnA-as*.

Table S3. Strains and primers used in this work.

Table S4. Diameters of halos cited in figure 7 and 8.

Table S5. Stalk length measurement by BacStalk software cited in figure 8 and S3.

Table S6. RNAseq Rockhopper data cited in figure S1A.

Table S7. Raw data of growth curves cited in Figure 2, 6, S8 and S9.

Table S8. Raw data of cell length and width measure obtained by MicrobeJ cited in figure 2, 7, 8, S3, S8 and S14.

**Figure S1.**
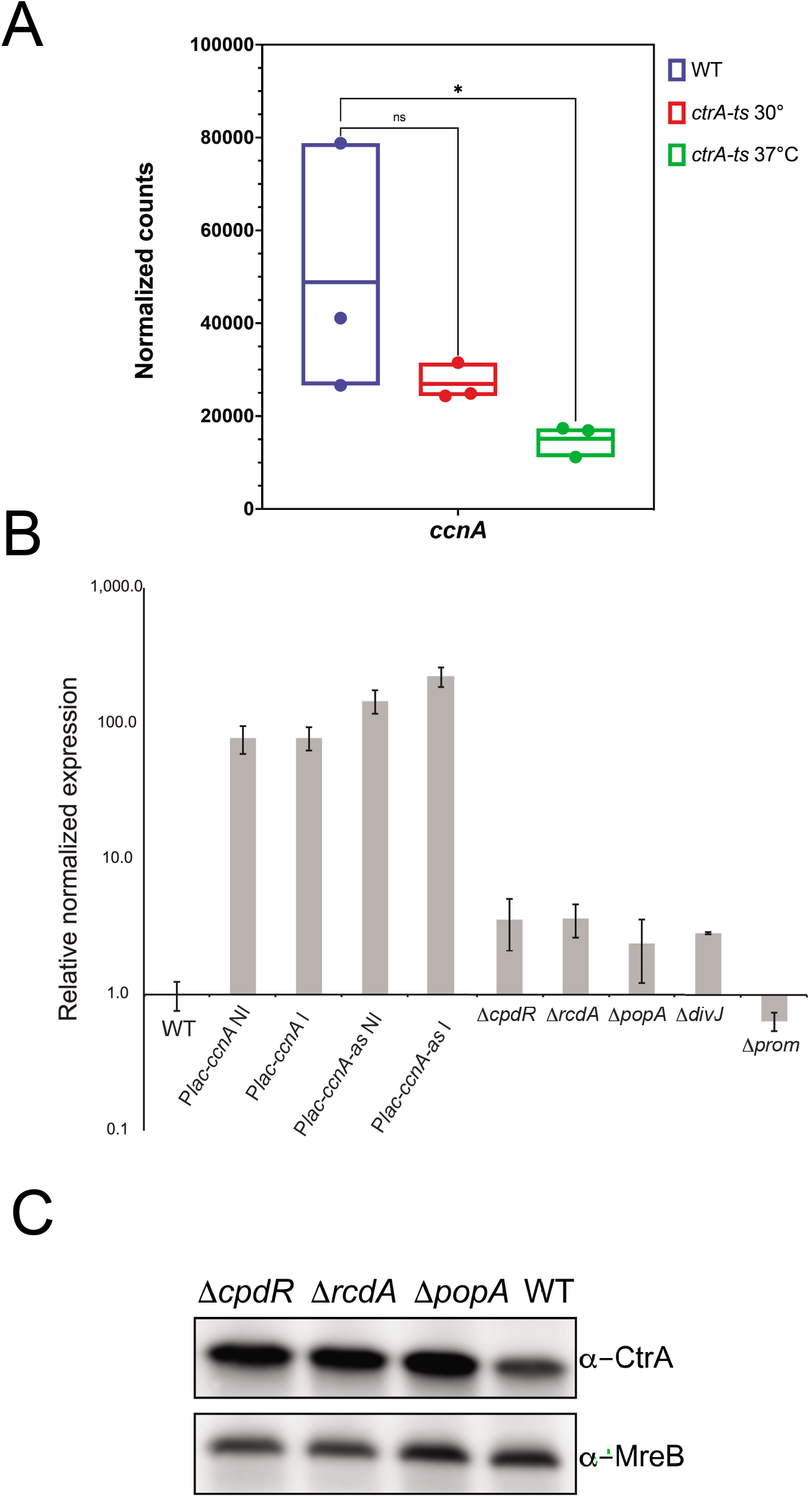
**A. Transcription levels of CcnA in a CtrA thermosensitive strain (*ctrAts*).** WT cells were grown in PYE until OD_600nm_= 0.6 and *ctrA-ts* cells were grown in PYE until OD_600nm_= 0.6 and kept for 1h at the permissive temperature 30°C or shifted 1h to the restrictive temperature 37°C. Total RNA was extracted in each condition and prepared for RNA sequencing. Data were analyzed using Rockhopper (McClure et al., 2013; Tjaden, 2015) with the following parameters: Strand specific, paired end, verbose output (generation of raw and normalized counts for each transcripts and RPKM value). Statistical analysis was performed using Kruskal-Wallis test with a Dunn’s multiple comparisons test (Prism GraphPad 9.1.2) and result are shown as mean (N=3) of normalized counts +/- SD, ns= difference not significant, * = p.val = 0,0225. Raw data are provided in Table S6. **B.** Expression level of CcnA in *C. crescentus* strains carrying *ccnA* and *ccnA-as* in a pSRK plasmid with or without induction by addition of IPTG 1mM 30min (related to figure 1 and S9) and in different backgrounds (WT, Δ*cpdR*, Δ*rcdA*, Δ*popA*, Δ*divJ* and Δ*prom*). Cells were grown in PYE at 30°C then, the expression of *ccnA* was determined by qRT-PCR and compared to 16S level. Results are presented as mean (N=3) +/- SD. **C.** WT *C. crescentus* cells, or *C. crescentus* cells deleted from the genes *cpdR* (Δ*cpdR*), *rcdA* (Δ*rcdA*) or *popA* (Δ*popA*), belonging to the proteolysis machinery ClpXP of *C. crescentus*, were grown in PYE at 30°C until OD_600nm_=0.6. Proteins were extracted and separated on a SDS-PAGE gel for Western blotting. CtrA and MreB (loading control) were revealed using specific polyclonal antibodies on nitrocellulose membranes.

**Figure S2.**
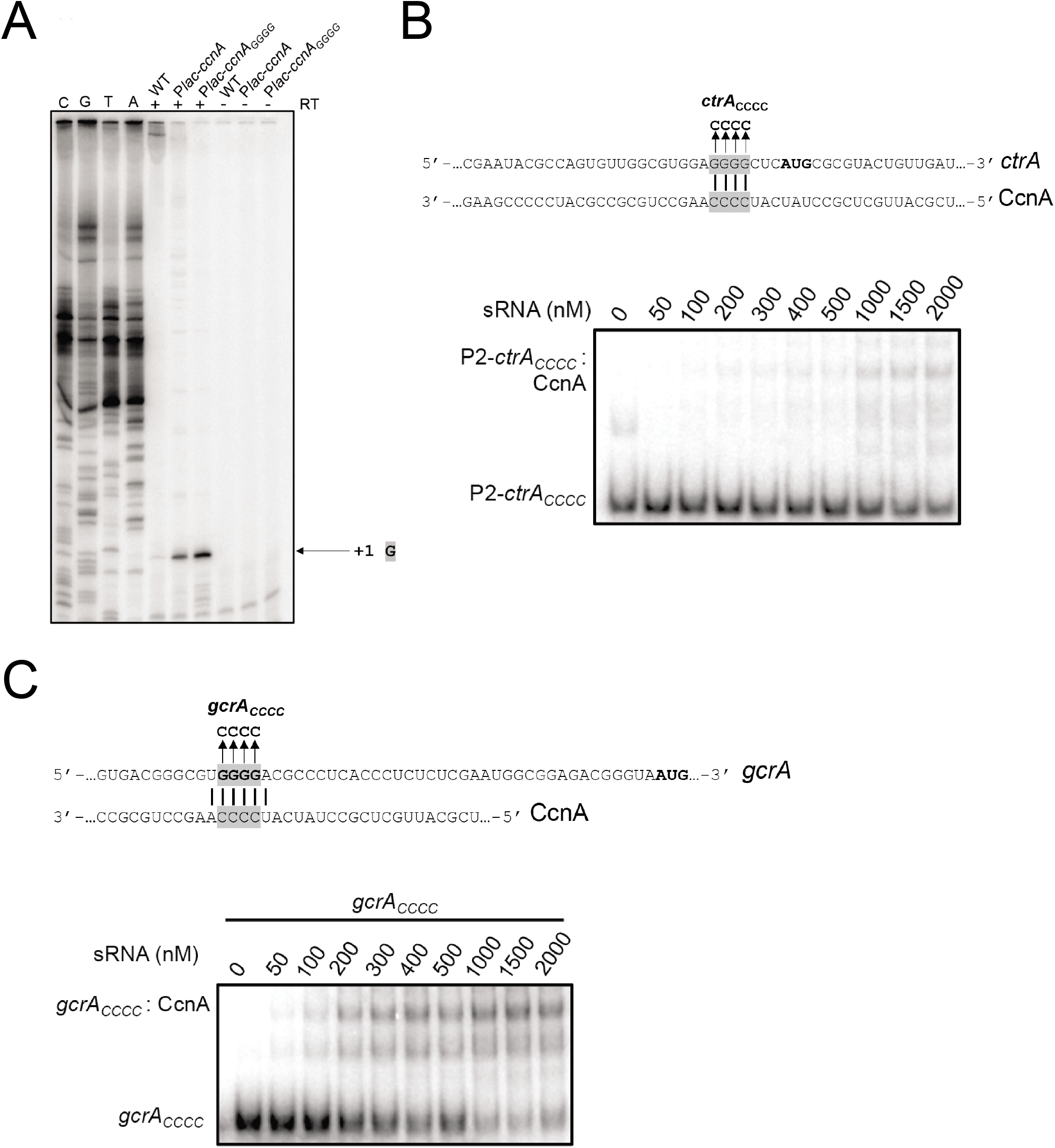
**A. Determination of the transcriptional +1 site of CcnA by primer extension related to figure 1 and 5.** Total RNA extracted from wild-type cells (WT) or containing P*lac-ccnA and* P*lac-ccnA-Loop A* were used with radiolabelled oligo. The reaction was done with (+) or without (-) reverse transcriptase. The same oligo was used for *ccnA* sequencing (GCAT). The +1 signal is represented by the arrow. **B. Mutation of the *ctrA* mRNA (*ctrA_CCCC_*) and ncRNA CcnA binding site.** Solid lines indicate CcnA binding sites on *ctrA*. Boxed gray text corresponds to the nucleotides mutated. The translation start codon is shown in bold. P2-*ctrA_CCCC_* (+143nt from the P2-*ctrA* promoter) 5nM RNA fragment was incubated with increasing concentration of CcnA (bottom). Data represent the mean of two independent experiments. **C. Mutation of the *gcrA* mRNA (*gcrA_CCCC_*) and ncRNA CcnA binding site**. Solid lines indicate CcnA binding sites on *gcrA*. Boxed gray text corresponds to the nucleotides mutated. *gcrA_CCCC_* 5nM RNA fragment was incubated with increasing concentration of CcnA (bottom). Data represent two independent experiments.

**Figure S3.**
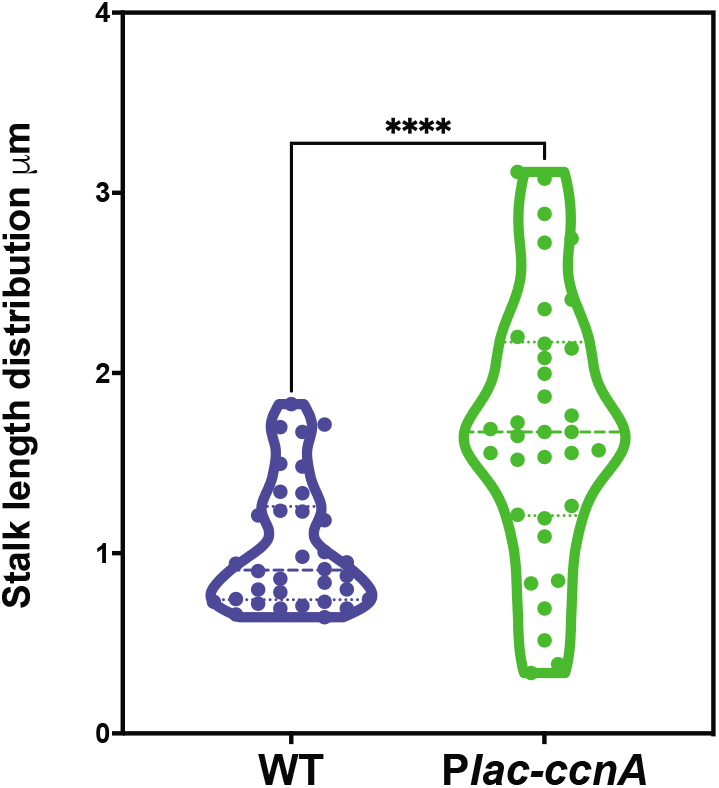
Violin plots of stalks length per cell for a WT + P*lac-ccnA* strain tested in Figure 2D compared to a WT *C. crescentus* used as a control for normal stalk length (same as 8D). Stalk length was measured by using BacStalk software (Hartmann et al., 2020). Statistical significance was determined using an unpaired t test with Welch’s correction for different SD. ****: p.val<0.0001. Raw data are in table S5.

**Figure S4.**
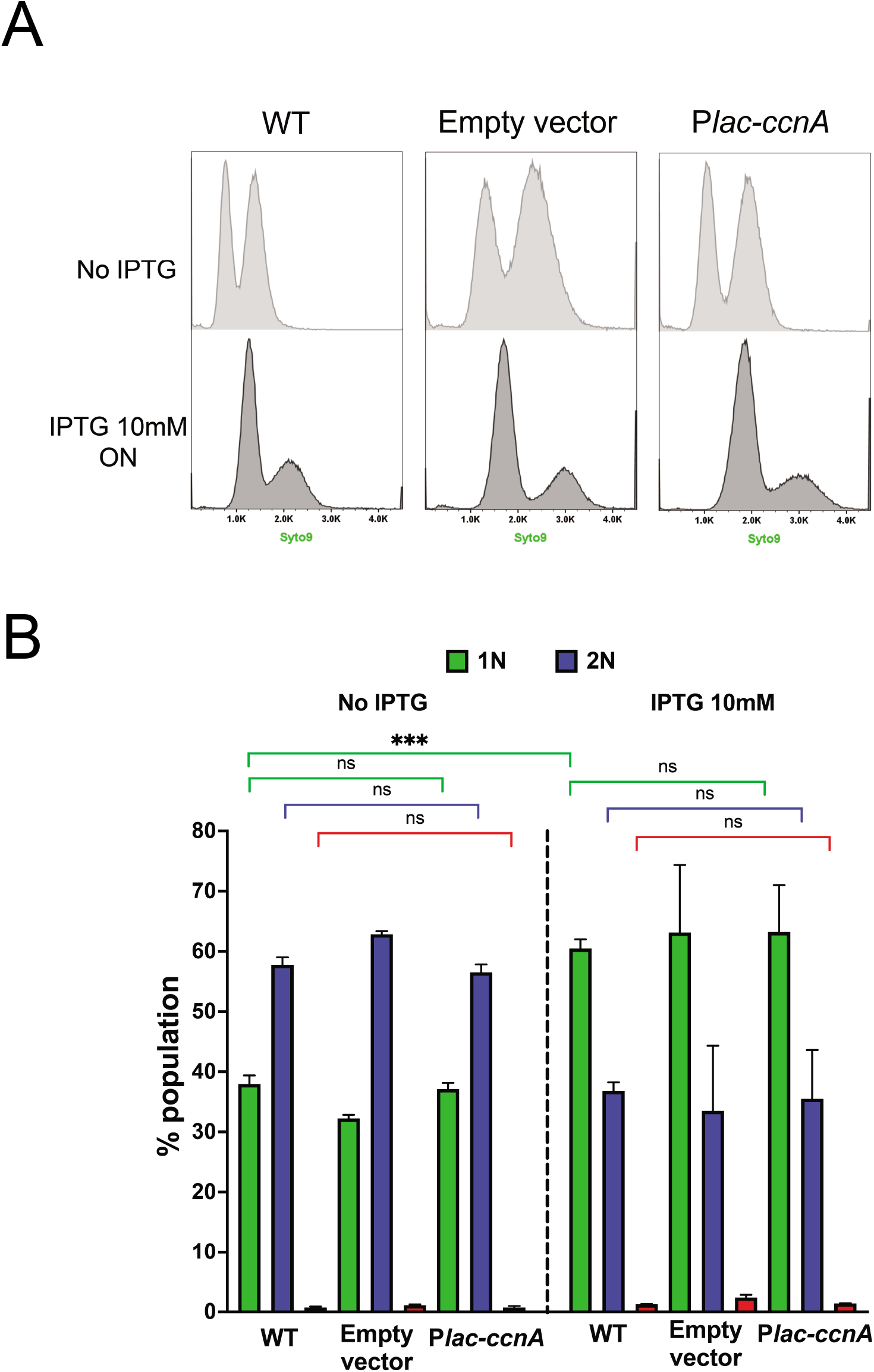
**A.** Flow cytometry profiles after SYTO 9 staining showing DNA content of WT cells, WT cells carrying either an empty pSRK (empty vector) or *ccnA* under the control of a P*lac* promoter (P*lac-ccnA*) grown in PYE until OD_600nm_ = 0.6. Then induction of P*lac-ccnA* was made overnight by addition of IPTG 10mM overnight. A total number of 300 000 particles were analysed per sample by flow cytometry. **B.** Proportions of cells harboring 1N, 2N and ≥3N DNA in the population were analyzed by gating the histograms in B). Data are representative of three biological replicates. Statistical analyses were carried out using ANOVA Tukey test. ns: difference not significant, *** : p.val. <0.001.

**Figure S5.**
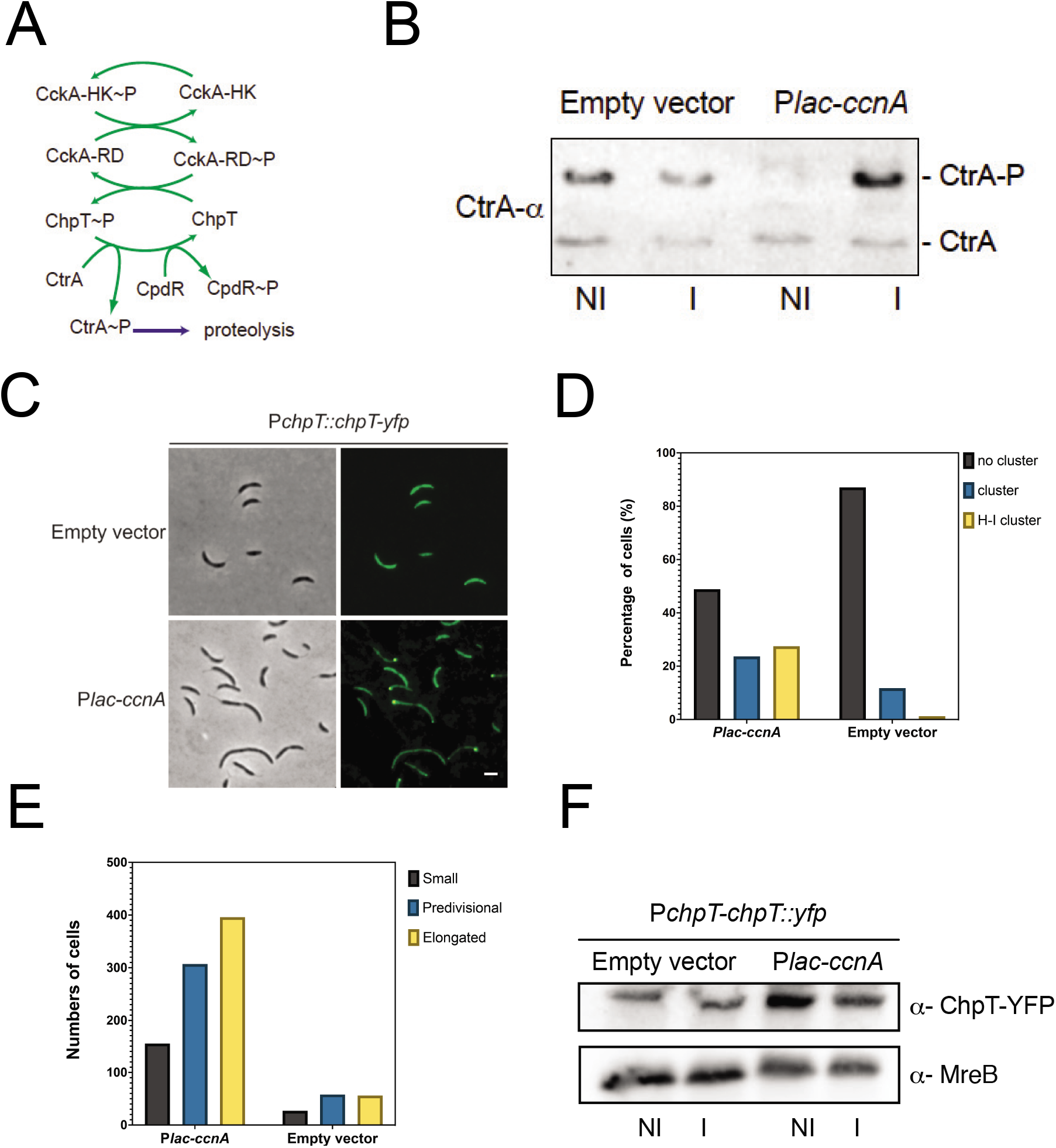
**A.** Schematic of the CckA-ChpT-CtrA phosphorelay. CckA is a hybrid histidine kinase, at the swarmer cell it acts as a kinase and at the stalked cell pole as a phosphatase. CckA autophosphorylates, then phosphorylates the histidine phosphotransferase ChpT that in return phosphorylates CtrA and CtrA proteolysis adapter protein CpdR. **B.** WT cells carrying either an empty pSRK (empty vector) or *ccnA* under the control of a P*lac* promoter (*Plac-ccnA*) were grown in PYE at 30°C until OD_600nm_ = 0.6. Then, the induction of P*lac-ccnA* was made by addition of IPTG 1mM 30min. Proteins were extracted and separated on a SDS-PAGE containing Phostag and Mn^2+^ to visualize CtrA phosphorylation level. CtrA was revealed using specific polyclonal antibodies on nitrocellulose membrane. **C.** Phase contrast and epifluorescence images of P*chpT-chpT::yfp* cells carrying *ccnA* (*Plac-ccnA*) or an empty pSRK (empty vector) grown in PYE at 30°C until OD_600nm_= 0.6. Scale bar= 2um. **D**. Histograms showing quantification of ChpT-YFP signal in P*chpT-chpT::yfp* cells carrying P*lac-ccnA* or an empty vector by using MicrobeJ. Results are shown as fraction of total cells with absence of cluster, presence of cluster and presence of high intensity cluster. At least 1000 cells were analyzed for each condition. **E.** Histograms showing morphologies typology of cells related to S5 C-D with detected cluster using MicrobeJ software (Ducret et al., 2016a). **F**. P*chpT-chpT::yfp* cells carrying either an empty vector or P*lac-ccnA* were grown in PYE 30°C until OD_600_ = 0.6. Then, the induction of P*lac-ccnA* was made by addition of IPTG 1mM 30min. Proteins were extracted and separated on a SDS-PAGE for western blotting. GFP and MreB (loading control) were revealed using specific polyclonal antibodies on nitrocellulose membrane.

**Figure S6.**
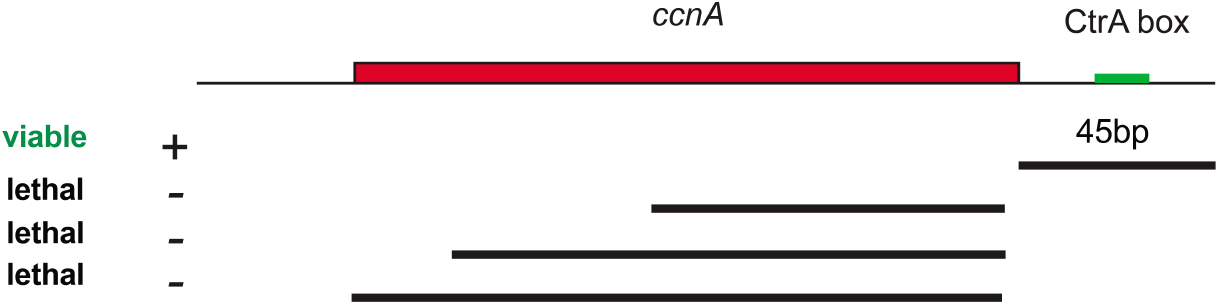
Schematic representation of the strategy used to study *ccnA* down-regulation phenotypes without lethality. Our only viable deletion obtained in this study was a 45bp long deletion comprising the CtrA box located within the promoter region of *ccnA*. Attempts to delete the *ccnA* gene sequence could not be constructed.

**Figure S7.**
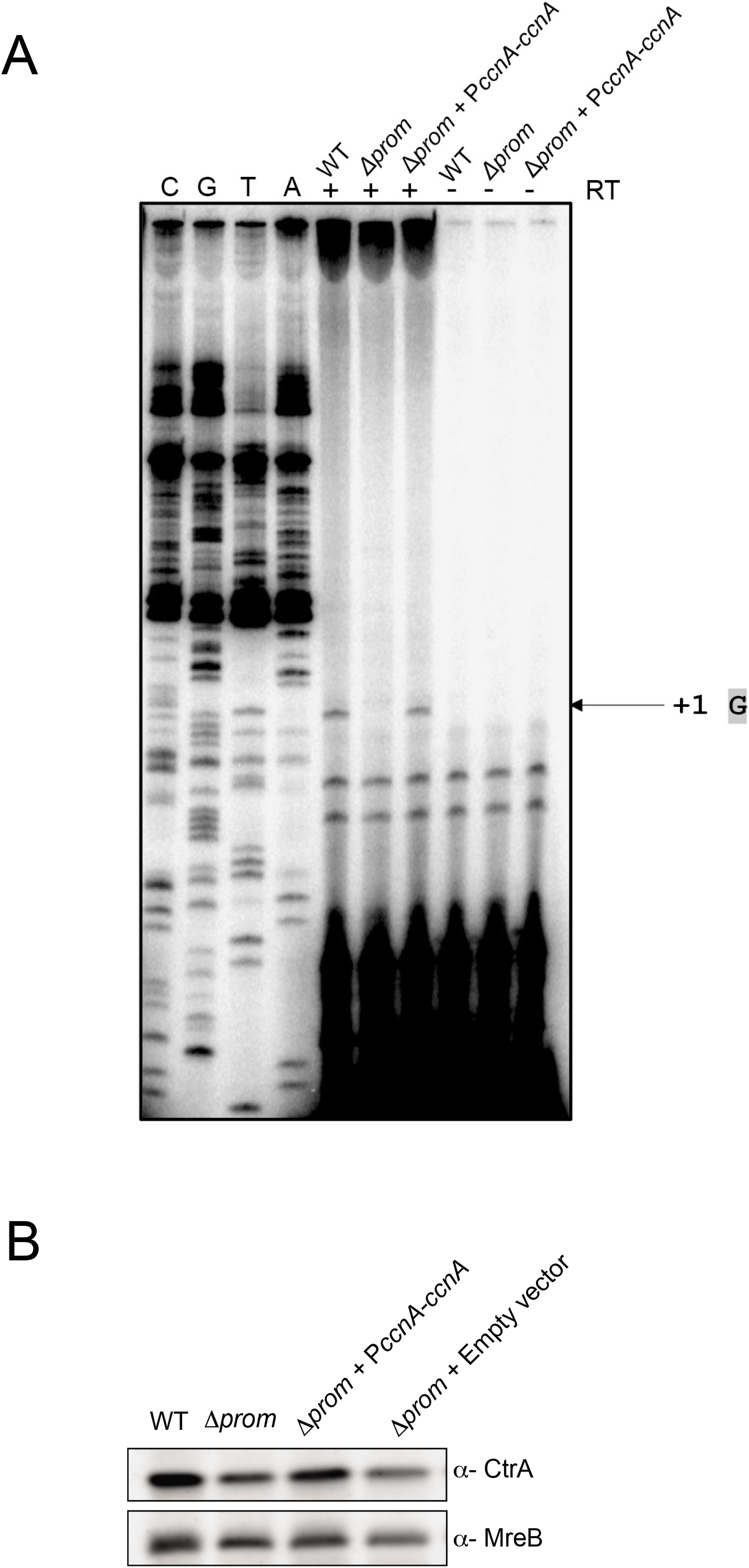
**A. Determination of the transcriptional +1 site of CcnA ncRNA by primer extension.** Total RNA extracted from wild-type cells (WT), deleted *ccnA* promoter (*Δprom*) and containing P*ccnA-ccnA* (Δ*prom* + P*ccnA-ccnA*) was used with radiolabelled oligo. The reaction was done with (+) or without (-) reverse transcriptase. The same oligo was used for *ccnA* sequencing (GCAT). The sequence is presented as the reverse complement. The +1 signal is represented by the arrow. **B.** WT cells, Δ*prom* cells, Δ*prom* cells carrying either an empty pMR10 low copy plasmid (Δ*prom + empty vector*) or harboring *ccnA* under the control of its own promoter (*Δprom* + P*ccnA-ccnA*) were grown in PYE at 30°C until OD_600nm_ = 0.6. Proteins were extracted and separated on a SDS-PAGE for western blotting. CtrA and MreB (loading control) were revealed using specific polyclonal antibodies on nitrocellulose membranes.

**Figure S8.**
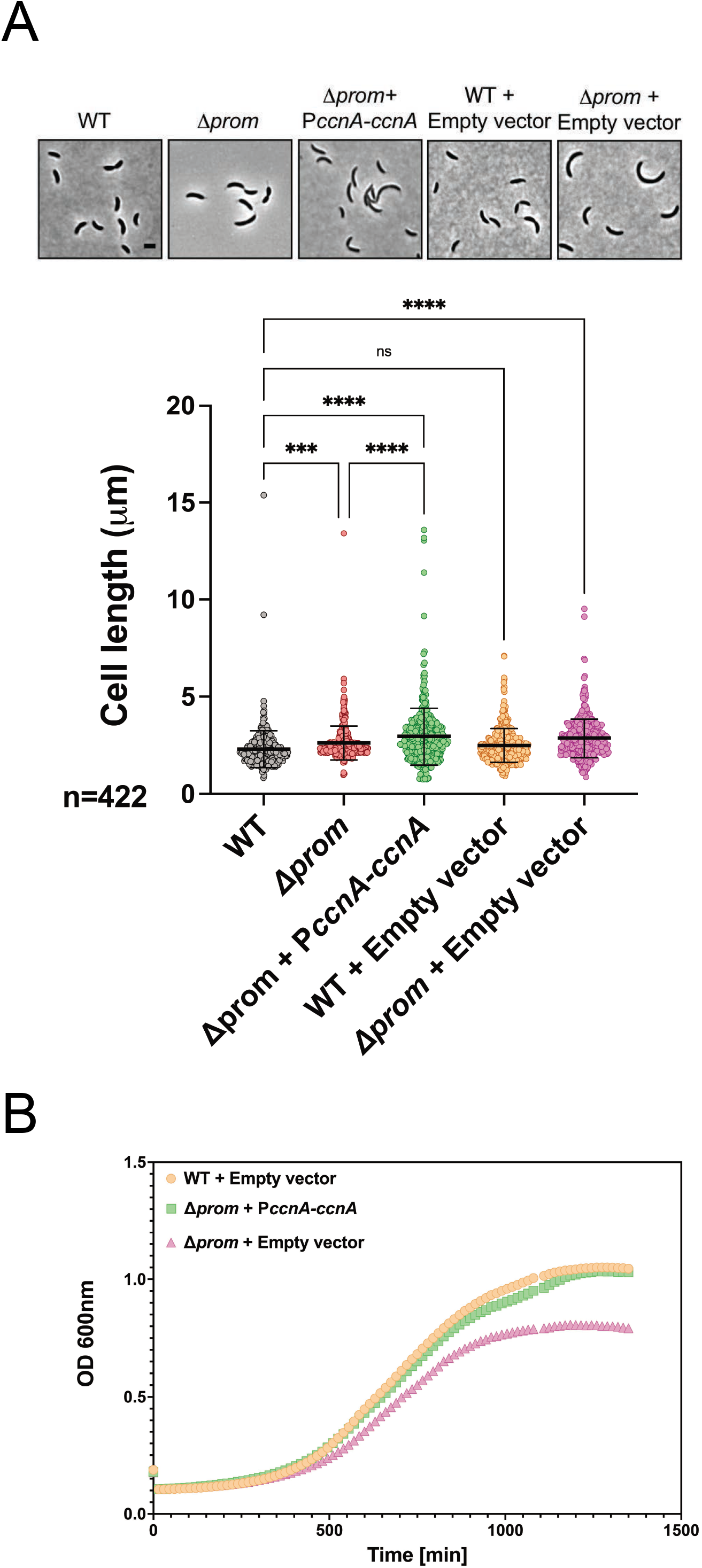
**A.** Phase contrast images of WT cells, Δ*prom* cells, Δ*prom* cells carrying *ccnA* under the control of its own promoter in a pMR10 low copy plasmid (Δ*prom* + P*ccnA-ccnA*), WT cells carrying an empty low copy plasmid pMR10 (WT + empty vector) or Δ*prom* cells carrying an empty low copy plasmid pMR10 (Δ*prom +* empty vector) grown overnight in PYE at 30°C until OD_600nm_= 0.6. Scale bar corresponds to 2 μm. Cells were analyzed using MicrobeJ (Ducret et al., 2016b) to assess cell length. 422 cells were analyzed. Statistical significance was determined using ANOVA with Tukey’s multiple comparisons test. ns: difference not significant *** : p.val= 0.0001 and ****: p.val <0.0001. Raw data are provided in Table S8. **B. Growth curves of Δ*prom* cells complemented with *ccnA***. WT cells carrying an empty low copy plasmid pMR10 (WT + empty vector), Δ*prom* cells carrying either *ccnA* under the control of its own promoter in a pMR10 low copy plasmid (Δ*prom* + P*ccnA-ccnA*) or an empty low copy plasmid pMR10 (Δ*prom +* empty vector) were grown overnight in PYE. 200μL of cells back-diluted from stationary phase cultures to an OD_600nm_= 0.02 were then grown on 96 wells in PYE at 30°C. Cell growth was monitored overnight with a Spark-TM at 30°C and a shaking (orbital) amplitude of 6 mm and a shaking (orbital) Frequency of 96 rpm. Results are shown as mean N=2 biological with 3 technical replicates per biological replicates. Raw Data are provided in Table S7.

**Figure S9.**
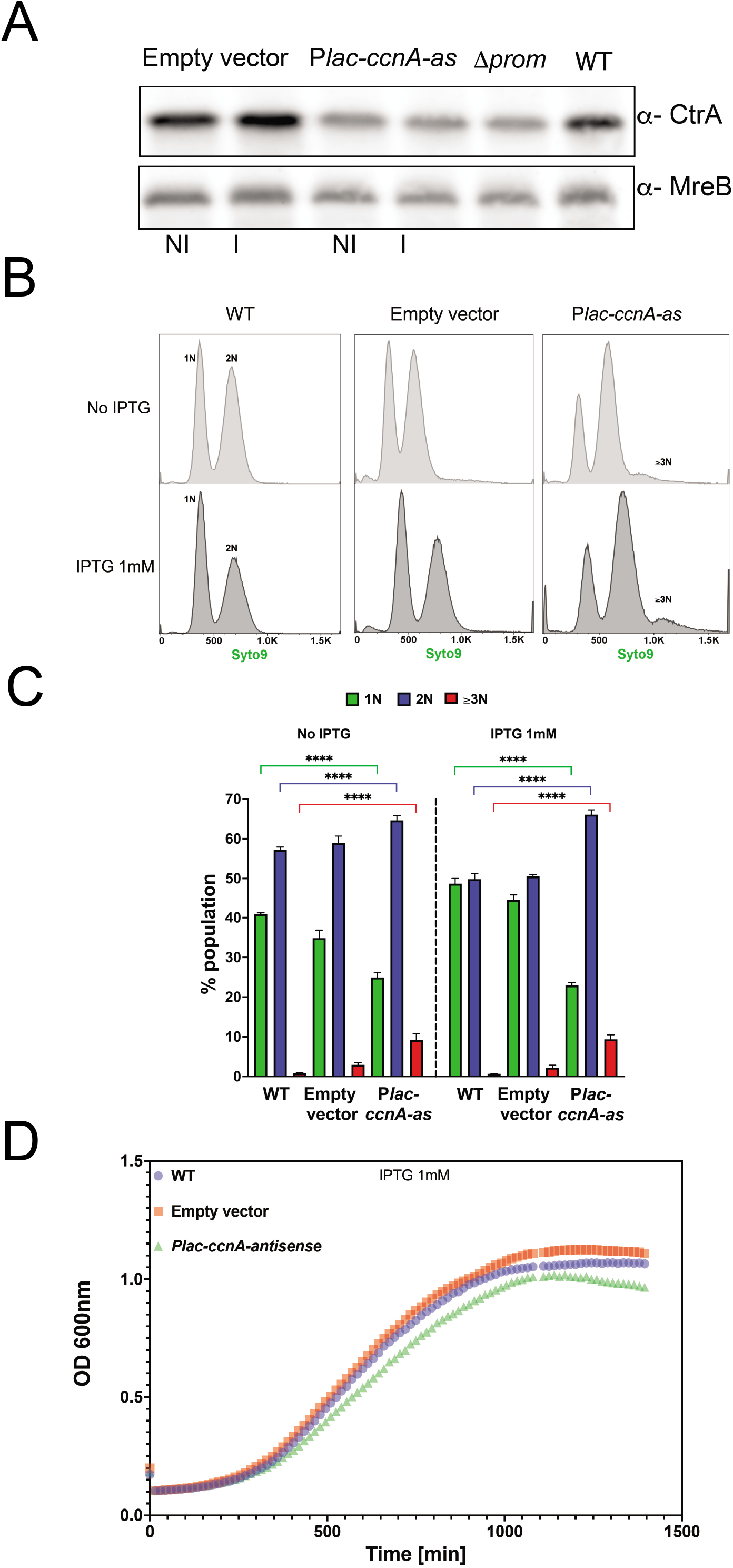
A. WT cells, WT cells carrying either an empty pSRK (empty vector) or *ccnA-antisense* under the control of a P*lac* promoter (P*lac-ccnA-as*) were grown in PYE at 30°C until OD_600nm_= 0.6. Then, the induction of P*lac-ccnA-as* was made by addition of IPTG 1mM 30min. As a control of induction WT cells carrying an empty vector were also incubated 30min in presence of IPTG 1mM and WT cells with no induction were used as a control. Δ*prom* cells were used as a comparison for CtrA levels. Proteins were extracted and separated on a SDS-PAGE gel for Western blotting. CtrA and MreB (loading control) proteins were revealed using specific polyclonal antibodies on nitrocellulose membranes. **B.** Flow cytometry profiles after SYTO 9 staining showing DNA content of WT cells, WT cells carrying either an empty pSRK (empty vector) or *ccnA-antisense* under the control of a *Plac* promoter (P*lac-ccnA-as*) grown in PYE until OD_600nm_ = 0.6. Then induction of P*lac-ccnA-as* was made by addition of IPTG 1mM 1h30min. A total number of 300 000 particles were analyzed per sample by flow cytometry. **C.** Proportions of cells harboring 1N, 2N and ≥3N DNA in the population were analyzed by gating the histograms in C). Data are representative of three biological replicates. Sstatistical analyses were carried out using ANOVA Tukey test. ****: p.val <0.0001. **D. Growth curves following the expression of CcnA antisense (P*lac-ccnA-as*).** WT cells and WT cells carrying either an empty pSRK (empty vector) or a pSRK with *ccnA* under the control of an inducible P*lac* promoter (P*lac-ccnA*) or P*lac-ccnA-as* were grown overnight in PYE without IPTG. 200μL of cells back-diluted from stationary phase cultures to an OD_600nm_= 0.02 were grown on 96 wells in PYE supplemented with 1mM IPTG. Cell growth was monitored overnight with a Spark-TM at 30°C and a shaking (orbital) amplitude of 6 mm and a shaking (orbital) frequency of 96 rpm. Doubling time of P*lac-ccnA-as* cells equals to 201.96 min +/- 11.66 with 1mM IPTG. Results are shown as mean N=3 biological with 3 technical replicates. Raw data are provided in Table S7.

**Figure S10.**
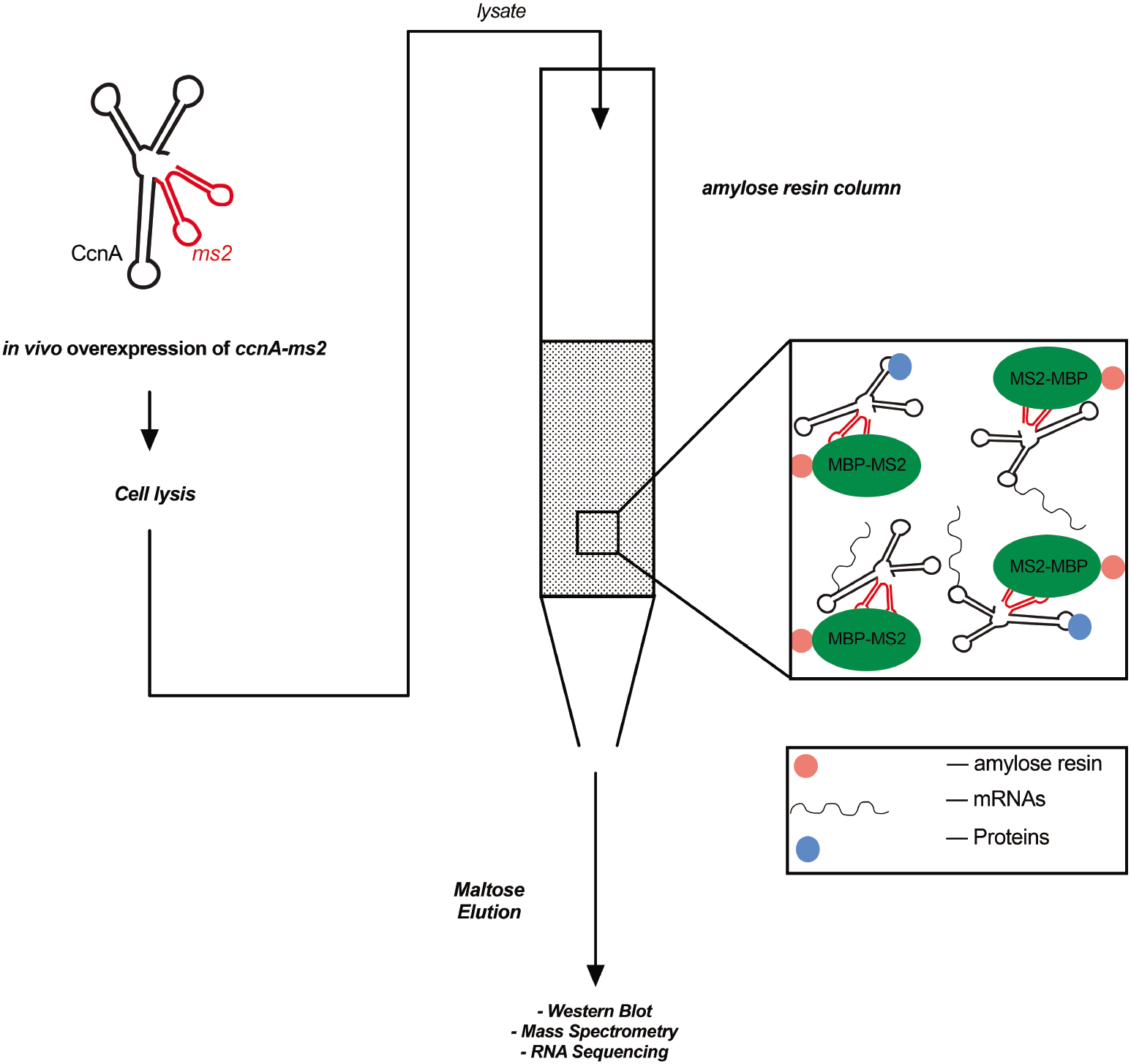
Schematic of the MAPS technique. MAPS consist on the fusion of a ncRNA of interest, here CcnA, with an RNA called MS2 (Lalaouna et al., 2017). In this technique the protein MS2 is used for its high affinity to the RNA MS2 from the bacteriophage MS2. For the experiment, the MS2 protein is fused to the maltose binding protein (MBP). The fused ncRNA is overexpressed *in vivo* prior to cell lysis. The soluble cellular (lysate) content is then transferred into a column containing an amylose resin added in order to fix the MS2-MBP fusion. Once the lysate is passed through the column, a solution of maltose is used to pull-down the MS2-CcnA complexes with RNAs or proteins interacting with CcnA. The identification and characterization of direct *in vivo* partners of CcnA is then made by Western blotting, Mass Spectrometry or RNA sequencing.

**Figure S11.**
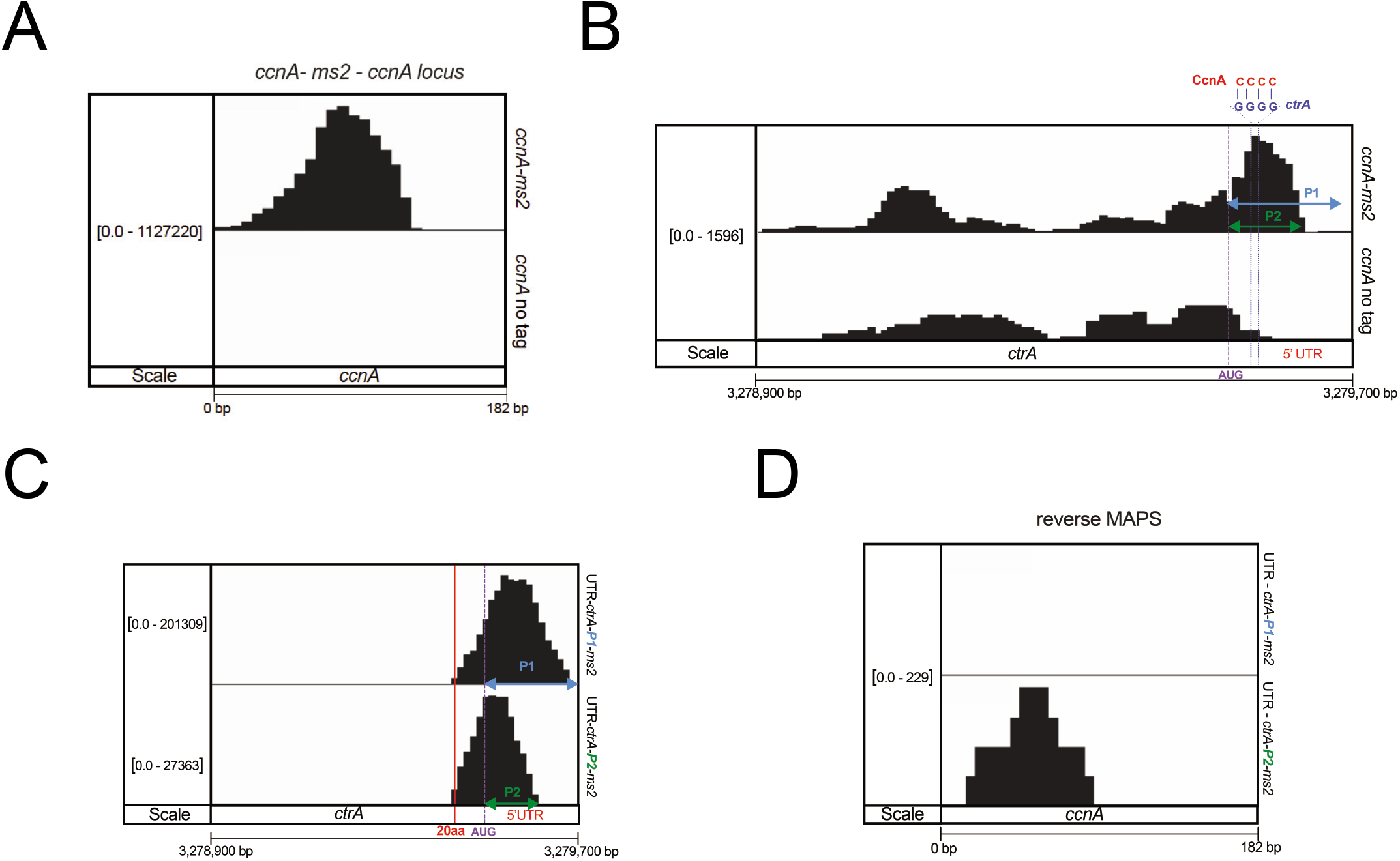
**A.** MAPS experiment was performed using an MS2-CcnA construct. Untagged CcnA was used as a control. Cells were grown in PYE at 30°C until OD_600nm_= 0.6 and harvested after induction of P*lac-ms2-ccnA* and P*lac-ccnA* by addition of IPTG 1mM 30min. After pull-down and RNA sequencing, data were normalized using RPKM method. Mapped reads of *ccnA* locus are visualized by using IGV. **B**. Same as (A) but here mapped reads of *ctrA* locus are visualized. The highest peak corresponds to a putative interaction site (CcnA in red; 5’UTR of *ctrA* mRNA in blue). **C.** Same as (A) but MAPS was performed using MS2-5’UTRs of *ctrA* generated by promoters *ctrA* P1 (UTR-*P1-ctrA-ms2*) or *ctrA* P2 (UTR-*P2-ctrA-ms2*). Mapped reads of P1 and P2 5’UTRs in the vicinity of *ctrA* mRNA were visualized. The red line shows reads corresponding to transcription until 20 amino acids of *ctrA*. **D**. Same as (C) but here mapped reads in the vicinity of *ccnA* were visualized by using IGV.

**Figure S12.**
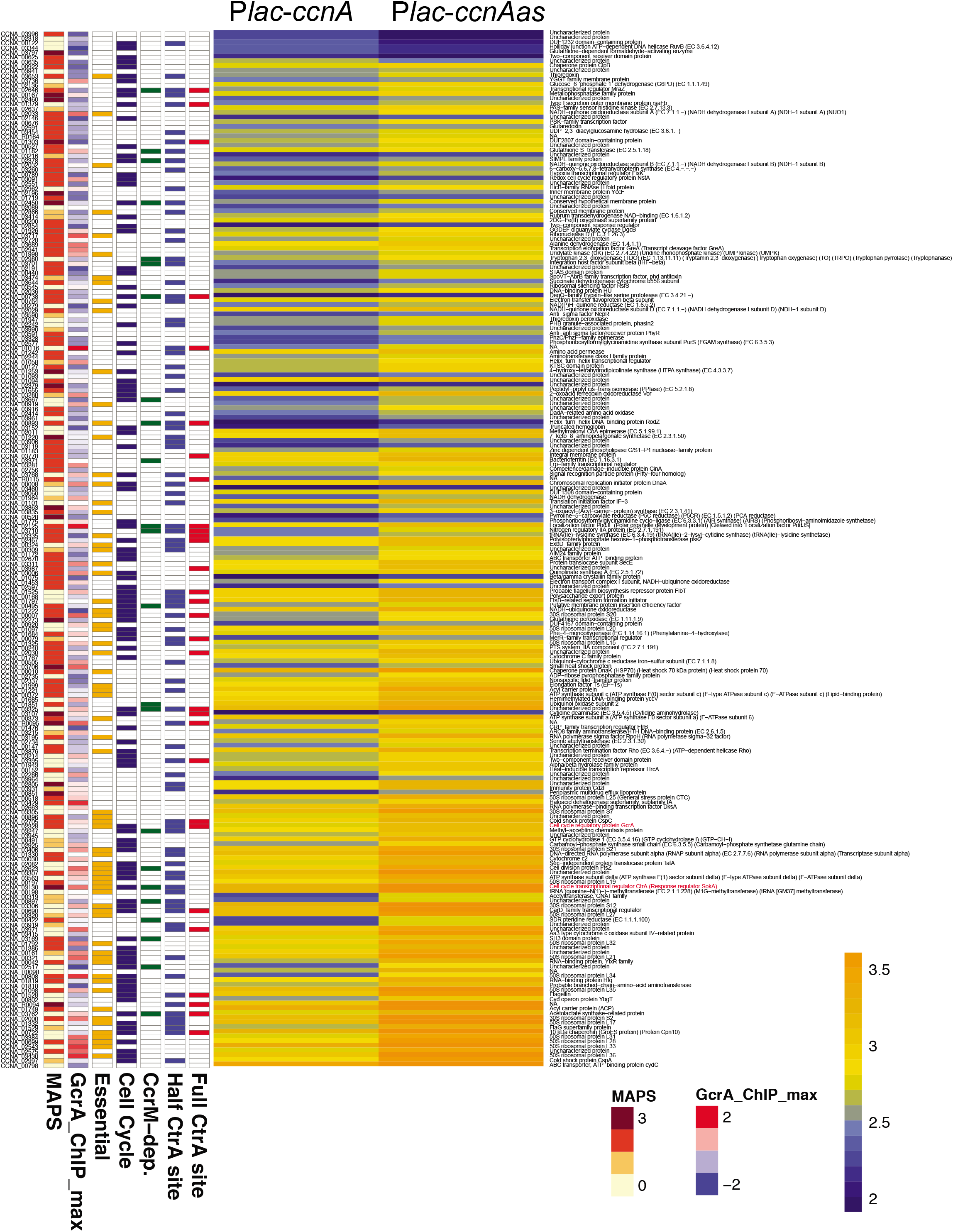
Characterization of the CcnA ncRNA *in vivo* targetome. MAPS experiment was performed using an MS2-CcnA construct. Untagged CcnA was used as a control. Cells were grown in PYE at 30°C until OD_600nm_= 0.6 and harvested after induction of P*lac-ms2-ccnA* and P*lac-ccnA* by addition of IPTG 1mM 30min. After pull-down and RNA sequencing reads were mapped to the indexed *C. crescentus* NA1000 genome (NC_011916) with Bowtie2 and mRNAs directly bound or not to CcnA were identified. In parallel, an additional RNA sequencing of transcriptome of cells carrying either P*lac-ccnA* or P*lac-ccnA-as* was performed in order to analyze expression profiles of CcnA targets. Detailed protocol for MAPS analysis is described in the material and methods section. Data are representative of three biological replicates. CtrA and GcrA are highlighted in red.

**Figure S13.**
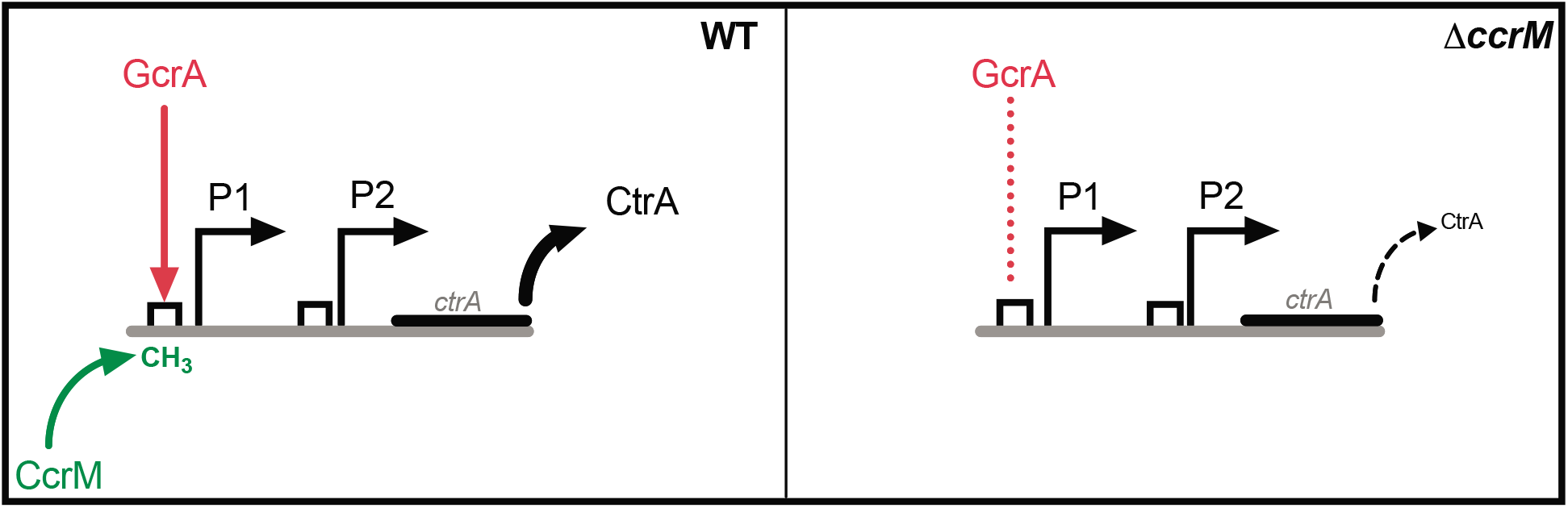
Schematic representation of the GcrA-CcrM epigenetic transcriptional regulation of *ctrA-P1*. The expression of *ctrA* depends on the GcrA-CcrM module. CcrM mediates the methylation of the P1 promoter of *ctrA*, recruiting GcrA which in turn activates the transcription of CtrA. In a Δ*ccrM* background the levels of CtrA are low. However, considering that the *ctrA P1* promoter is sigma70 dependent, transcription of *ctrA* will keeps a basal low expression.

**Figure S14.**
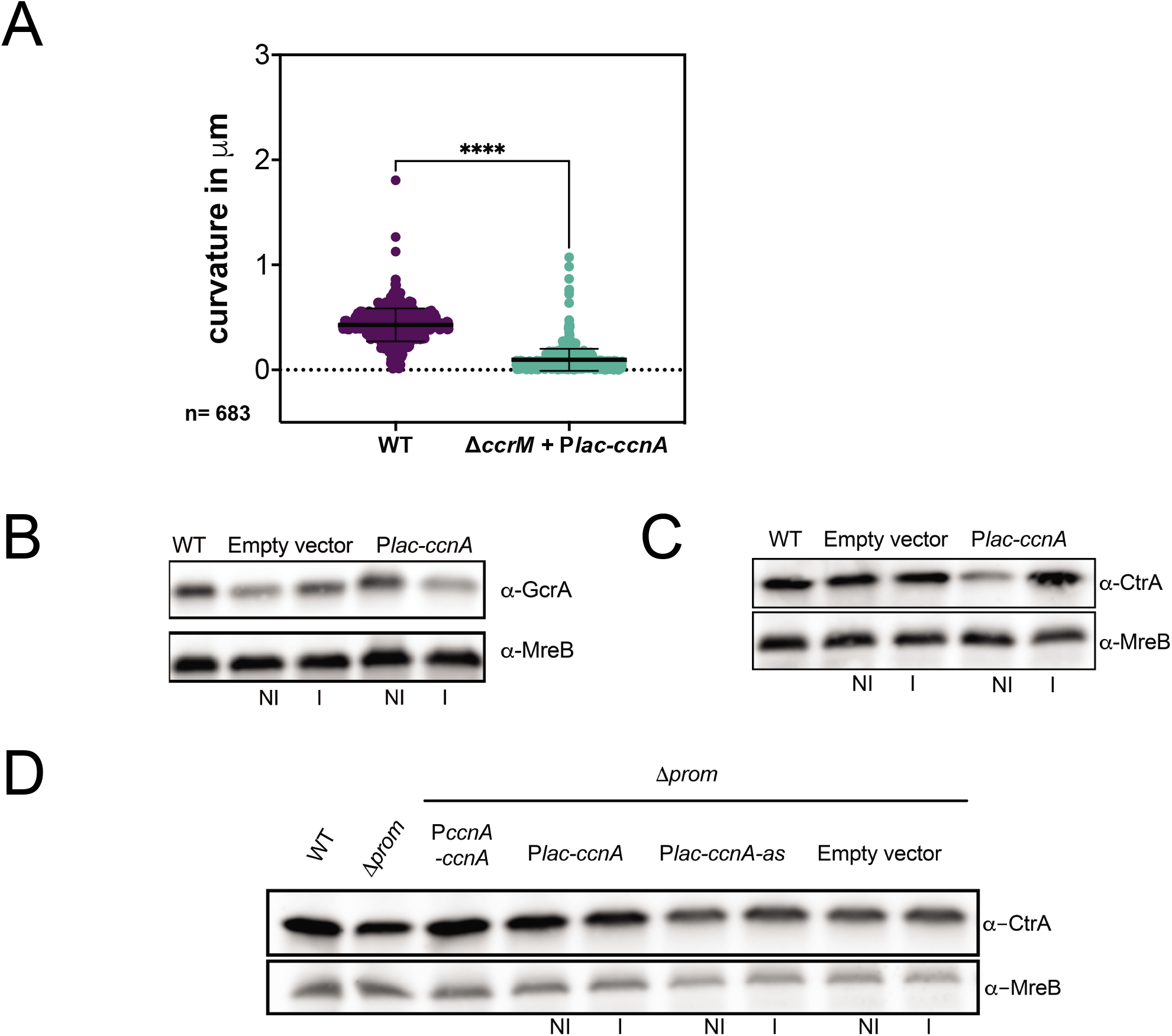
**A.** Cell curvature of WT cells and Δ*ccrM+* P*lac-ccnA* from (6A) was determined using MicrobeJ (Ducret et al., 2016a). 683 cells were analyzed for each condition and statistical significance was determined using an unpaired t test. ****: p.val <0.0001. The *ΔccrM* and *ΔccrM* + empty vector cells were not considered for this analysis because the filamentous phenotypes observed in these two mutants result in cell curvature artifacts measurements that are not consistent with the MicrobeJ analysis. Raw data are provided in Table S8. **B.** WT cells, WT cells carrying an empty vector or *Plac-ccnA* were grown in PYE at 30°C until OD_600nm_ = 0.6. Then, induction of P*lac-ccnA* was made by addition of IPTG 1mM 30min. As a control of induction WT cells carrying an empty vector were also incubated 30min in presence of IPTG 1mM and WT cells with no induction were used as a control. (NI= no IPTG) and (I= IPTG). Proteins were extracted and separated on a SDS-PAGE gel for Western blotting. GcrA and MreB (loading control) proteins were revealed using specific polyclonal antibodies on nitrocellulose membranes. **C.** Same as (B) but here proteins from a biological replicate of P*lac-ccnA* cells were extracted and separated on a SDS-PAGE gel for Western blotting. CtrA and MreB (loading control) proteins were revealed using specific polyclonal antibodies on nitrocellulose membranes. **D.** WT cells, Δ*prom* cells, Δ*prom* cells carrying either a pMR10 low copy plasmid harboring *ccnA* under the control of its own promoter (Δ*prom* + P*ccnA-ccnA*) or *ccnA*, its antisense under the control of a P*lac* promoter (Δ*prom*+ P*lac-ccnA*, Δ*prom* + P*lac-ccnA-as*) and an empty pSRK used as a control (*Δprom* + empty vector) were grown in PYE at 30°C until OD_600nm_= 0.6. For Δ*prom*+ P*lac-ccnA*, Δ*prom* + P*lac-ccnA-as* cells, expression of *ccnA* or its antisense was made by addition of IPTG 1mM 30min. As a control Δ*prom* cells carrying the empty pSRK were also incubated 30min in presence of IPTG 1mM 30min. Proteins were extracted and separated on a SDS-PAGE gel for Western blotting. CtrA and MreB (loading control) proteins were revealed using specific polyclonal antibodies on nitrocellulose membranes

**Figure S15.**
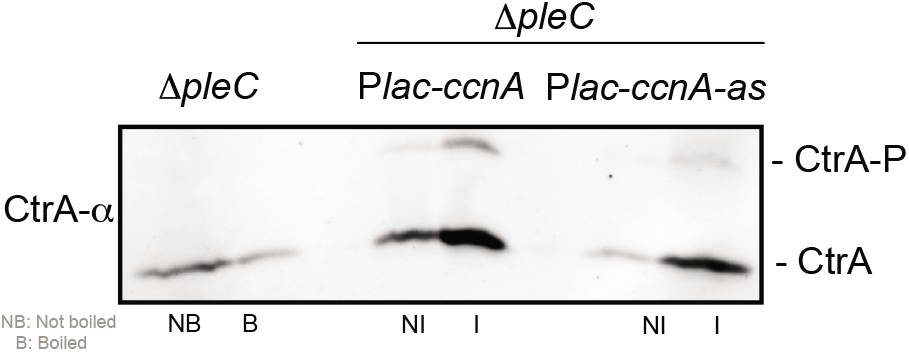
Δ*pleC+Plac-ccnA* or Δ*pleC*+ P*lac-ccnA-as* were grown in PYE at 30°C until OD_600nm_ = 0.6. Then, induction of *Plac-ccnA* and *Plac-ccnA-as* was made by addition of IPTG 1mM 30min. In parallel, as a control, Δ*pleC* cells were grown in PYE at 30°C until OD_600nm_= 0.6 and harvested. Proteins were extracted and separated on a SDS-PAGE gel containing Phostag and Mn^2+^ to visualize CtrA phosphorylation level. As a Phostag control, an additional sample of Δ*pleC* cells pellet was boiled 10min in order to discriminate the migration on the gel of the CtrA phosphorylated band from the non-phosphorylated band. CtrA was revealed using specific polyclonal antibodies on nitrocellulose membranes.

**Figure S16.**
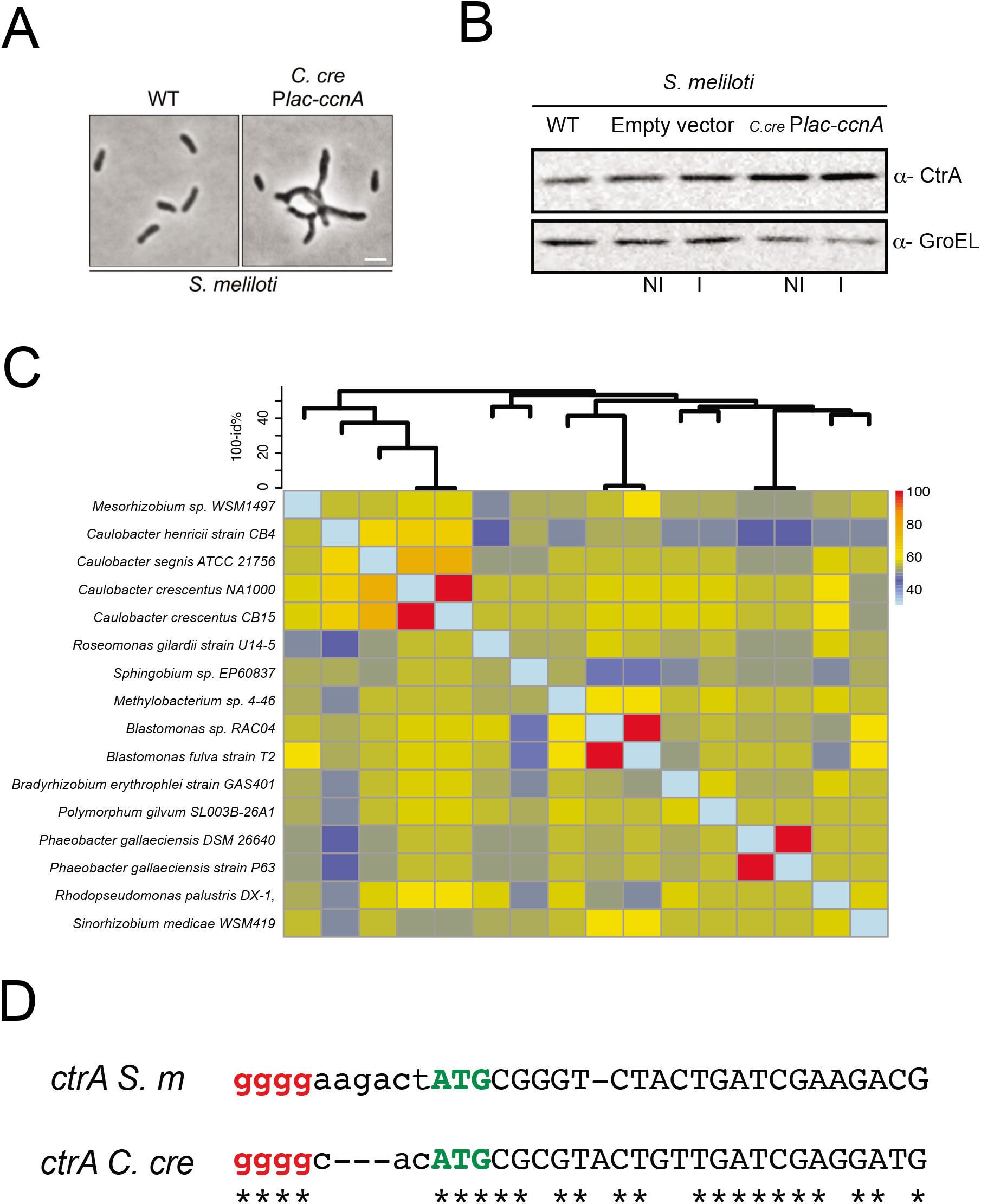
**A.** Phase contrast images of *S. meliloti* WT cells, carrying either an empty pSRK (Empty vector) or *ccnA* under the control of a P*lac* promoter (P*lac-ccnA*) from *C. crescentus* were grown in TY at 30°C until OD_600nm_ = 0.6. **B.** WT *S. meliloti* carrying either an empty vector or P*lac-ccnA* were grown in TY at 30°C until OD_600nm_= 0.6. Then, the induction of P*lac-ccnA* was made by addition of IPTG 1mM 30min. Proteins were extracted and separated on a SDS-PAGE gel for Western blotting. CtrA and GroEL (loading control) proteins were revealed using specific polyclonal antibodies on a nitrocellulose membrane. **C.** Putative homologs of *ccnA* in the class *Alphaproteobacteria*. Research of homologs was performed using the online sRNA Homolog Finder GlassGo (Lott et al., 2018b) using *C. crescentus ccnA* sequence as query. The heatmap contains identity percentages shared by CcnA homologs in different species and was then transformed into a distance matrix to build the dendrogram on the top. Comparisons were done in pairs because a multiple alignment of all CcnA homologs contains too many gaps. **D**. ClustalOmega (Madeira et al., 2019; Sievers et al., 2011) alignment of UTRs of *ctrA* from *S. meliloti* and *C. crescentus* starting from GGGG (red) motif near the start codon ATG (green) until nucleotide +25. Clustal Omega was used with default parameters for RNA. “*” represent a conserved nucleotide between the two sequences.

## Author Contribution

WB and EGB conceived the experiments and prepared the manuscript. All authors revised and edited the manuscript. KP performed Probing, EMSA and PE. DL and WB performed MAPS. WB performed RNAseq. WB ad MD performed microscopy analysis. WB performed synchronization and WB and GB performed Flow cytometry experiments and analysis. NB, OV participated in the cloning and Western blotting, respectively.

## Financial Disclosure

This research was funded by the Agence Nationale Recherche (ANR; ANR-17-CE20-0011-01) to EGB. The funders had no role in study design, data collection and analysis, decision to publish, or preparation of the manuscript.

## Notes

### Competing Interest Statement

The authors have declared no competing interest.

## References

1. Afgan, E., Baker, D., van den Beek, M., Blankenberg, D., Bouvier, D., Cech, M., Chilton, J., Clements, D., Coraor, N., Eberhard, C., et al. (2016). The Galaxy platform for accessible, reproducible and collaborative biomedical analyses: 2016 update. Nucleic Acids Res. 44, W3–W10.

2. Anders, S., Pyl, P.T., and Huber, W. (2015). HTSeq-A Python framework to work with high-throughput sequencing data. Bioinformatics 31, 166–169.

3. Beroual, W., Brilli, M., and Biondi, E.G. (2018). Non-coding RNAs Potentially Controlling Cell Cycle in the Model Caulobacter crescentus: A Bioinformatic Approach. Front. Genet. 9, 164.

4. Biondi, E.G., Reisinger, S.J., Skerker, J.M., Arif, M., Perchuk, B.S., Ryan, K.R., and Laub, M.T. (2006a). Regulation of the bacterial cell cycle by an integrated genetic circuit. Nature 444, 899–904.

5. Biondi, E.G., Skerker, J.M., Arif, M., Prasol, M.S., Perchuk, B.S., and Laub, M.T. (2006b). A phosphorelay system controls stalk biogenesis during cell cycle progression in Caulobacter crescentus. Mol. Microbiol. 59, 386–401.

6. Brilli, M., Fondi, M., Fani, R., Mengoni, A., Ferri, L., Bazzicalupo, M., and Biondi, E.G. (2010a). The diversity and evolution of cell cycle regulation in alpha-proteobacteria: a comparative genomic analysis. BMC Syst. Biol. 4, 52.

7. Brilli, M., Fondi, M., Fani, R., Mengoni, A., Ferri, L., Bazzicalupo, M., and Biondi, E.G.E.G. (2010b). The diversity and evolution of cell cycle regulation in alpha-proteobacteria: A comparative genomic analysis. BMC Syst. Biol. 4.

8. Chen, Y.E., Tropini, C., Jonas, K., Tsokos, C.G., Huang, K.C., and Laub, M.T. (2011). Spatial gradient of protein phosphorylation underlies replicative asymmetry in a bacterium. Proc. Natl. Acad. Sci. U. S. A. 108, 1052–1057.

9. Christen, B., Abeliuk, E., Collier, J.M., Kalogeraki, V.S., Passarelli, B., Coller, J.A., Fero, M.J., McAdams, H.H., and Shapiro, L. (2011a). The essential genome of a bacterium. Mol. Syst. Biol. 7, 528.

10. Christen, B., Abeliuk, E., Collier, J.M., Kalogeraki, V.S., Passarelli, B., Coller, J.A., Fero, M.J., Mcadams, H.H., and Shapiro, L. (2011b). The essential genome of a bacterium. Mol. Syst. Biol. 7, 1–7.

11. Collier, J. (2012). Regulation of chromosomal replication in Caulobacter crescentus. Plasmid 67, 76–87.

12. Collier, J., Murray, S.R., and Shapiro, L. (2006). DnaA couples DNA replication and the expression of two cell cycle master regulators. EMBO J. 25, 346–356.

13. Collier, J., McAdams, H.H., and Shapiro, L. (2007). A DNA methylation ratchet governs progression through a bacterial cell cycle. Proc. Natl. Acad. Sci. U. S. A. 104, 17111–17116.

14. Coppine, J., Kaczmarczyk, A., Petit, K., Brochier, T., Jenal, U., and Hallez, R. (2020). Regulation of Bacterial Cell Cycle Progression by Redundant Phosphatases. J. Bacteriol. 202, e00345–20.

15. Delaby, M., Panis, G., and Viollier, P.H. (2019). Bacterial cell cycle and growth phase switch by the essential transcriptional regulator CtrA. Nucleic Acids Res. 47, 10628–10644.

16. Ducret, A., Quardokus, E.M., and Brun, Y.V. (2016a). MicrobeJ, a tool for high throughput bacterial cell detection and quantitative analysis. Nat. Microbiol. 1, 16077.

17. Ducret, A., Quardokus, E.M., and Brun, Y.V. (2016b). MicrobeJ, a tool for high throughput bacterial cell detection and quantitative analysis. Nat. Microbiol. 1, 16077.

18. Dutta, T., and Srivastava, S. (2018). Small RNA-mediated regulation in bacteria: A growing palette of diverse mechanisms. Gene.

19. Fang, G., Passalacqua, K.D., Hocking, J., Llopis, P.M., Gerstein, M., Bergman, N.H., and Jacobs-Wagner, C. (2013a). Transcriptomic and phylogenetic analysis of a bacterial cell cycle reveals strong associations between gene co-expression and evolution. BMC Genomics 14, 450.

20. Fang, G., Passalacqua, K.D., Hocking, J., Llopis, P.M., Gerstein, M.B., Bergman, N.H., Jacobs-Wagner, C., Nicholas H Bergman, and Jacobs-Wagner, C. (2013b). Transcriptomic and phylogenetic analysis of a bacterial cell cycle reveals strong associations between gene co-expression and evolution. BMC Genomics 14, 450.

21. Fioravanti, A., Fumeaux, C., Mohapatra, S.S., Bompard, C., Brilli, M., Frandi, A., Castric, V., Villeret, V., Viollier, P.H., and Biondi, E.G. (2013). DNA Binding of the Cell Cycle Transcriptional Regulator GcrA Depends on N6-Adenosine Methylation in Caulobacter crescentus and Other Alphaproteobacteria. PLoS Genet. 9, e1003541.

22. Frandi, A., and Collier, J. (2019). Multilayered control of chromosome replication in Caulobacter crescentus. Biochem. Soc. Trans. 47, 187–196.

23. Fröhlich, K.S., Förstner, K.U., and Gitai, Z. (2018). Post-transcriptional gene regulation by an Hfq-independent small RNA in Caulobacter crescentus. Nucleic Acids Res. 46, 10969–10982.

24. Fumeaux, C., Radhakrishnan, S.K., Ardissone, S., Théraulaz, L., Frandi, A., Martins, D., Nesper, J., Abel, S., Jenal, U., and Viollier, P.H. (2014). Cell cycle transition from S-phase to G1 in Caulobacter is mediated by ancestral virulence regulators. Nat. Commun. 5, 4081.

25. Gonzalez, D., and Collier, J. (2013). DNA methylation by CcrM activates the transcription of two genes required for the division of Caulobacter crescentus. Mol. Microbiol. 88, 203–218.

26. Gonzalez, D., Kozdon, J.B., Mcadams, H.H., Shapiro, L., and Collier, J. (2014). The functions of DNA methylation by CcrM in Caulobacter crescentus: A global approach. Nucleic Acids Res. 42, 3720–3735.

27. Gora, K.G., Tsokos, C.G., Chen, Y.E., Srinivasan, B.S., Perchuk, B.S., and Laub, M.T. (2010). A cell-type-specific protein-protein interaction modulates transcriptional activity of a master regulator in Caulobacter crescentus. Mol. Cell 39, 455–467.

28. Gora, K.G., Cantin, A., Wohlever, M., Joshi, K.K., Perchuk, B.S., Chien, P., and Laub, M.T. (2013). Regulated proteolysis of a transcription factor complex is critical to cell cycle progression in Caulobacter crescentus. Mol. Microbiol. 87, 1277–1289.

29. Greene, S.E., Brilli, M., Biondi, E.G., and Komeili, A. (2012). Analysis of the CtrA pathway in Magnetospirillum reveals an ancestral role in motility in alphaproteobacteria. J. Bacteriol. 194, 2973–2986.

30. Haakonsen, D.L., Yuan, A.H., and Laub, M.T. (2015a). The bacterial cell cycle regulator GcrA is a σ70 cofactor that drives gene expression from a subset of methylated promoters. Genes Dev. 29, 2272–2286.

31. Haakonsen, D.L., Yuan, A.H., and Laub, M.T. (2015b). The bacterial cell cycle regulator GcrA is a $σ$ 70 cofactor that drives gene expression from a subset of methylated promoters. Genes Dev. 29, 2272–2286.

32. Hartmann, R., Teeseling, M.C.F. van, Thanbichler, M., and Drescher, K. (2020). BacStalk: A comprehensive and interactive image analysis software tool for bacterial cell biology. Mol. Microbiol. 114, 140–150.

33. Hartmann, R., Teeseling, M.C.F. van, Thanbichler, M., and Drescher, K. BacStalk: A comprehensive and interactive image analysis software tool for bacterial cell biology. Mol. Microbiol. n/a.

34. Holtzendorff, J., Hung, D., Brende, P., Reisenauer, A., Viollier, P.H., McAdams, H.H., and Shapiro, L. (2004). Oscillating global regulators control the genetic circuit driving a bacterial cell cycle. Science 304, 983–987.

35. Jacobs, C., Ausmees, N., Cordwell, S.J., Shapiro, L., and Laub, M.T. (2003). Functions of the CckA histidine kinase in Caulobacter cell cycle control. Mol. Microbiol. 47, 1279–1290.

36. Jagodnik, J., Chiaruttini, C., and Guillier, M. (2017). Stem-Loop Structures within mRNA Coding Sequences Activate Translation Initiation and Mediate Control by Small Regulatory RNAs. Mol. Cell 68, 158–170.e3.

37. Joshi, K.K., Bergé, M., Radhakrishnan, S.K., Viollier, P.H., and Chien, P. (2015). An Adaptor Hierarchy Regulates Proteolysis during a Bacterial Cell Cycle. Cell 163, 419–431.

38. Kaczmarczyk, A., Hempel, A.M., von Arx, C., Böhm, R., Dubey, B.N., Nesper, J., Schirmer, T., Hiller, S., and Jenal, U. (2020). Precise timing of transcription by c-di-GMP coordinates cell cycle and morphogenesis in Caulobacter. Nat. Commun. 11, 816.

39. Keiler, K.C., and Shapiro, L. (2003). tmRNA Is Required for Correct Timing of DNA Replication in Caulobacter crescentus. J. Bacteriol. 185, 573–580.

40. Khan, S.R., Gaines, J., Roop, R.M., 2nd, and Farrand, S.K. (2008). Broad-host-range expression vectors with tightly regulated promoters and their use to examine the influence of TraR and TraM expression on Ti plasmid quorum sensing. Appl. Environ. Microbiol. 74, 5053–5062.

41. Lalaouna, D., Carrier, M.-C., Semsey, S., Brouard, J.-S., Wang, J., Wade, J.T., and Massé, E. (2015). A 3’ external transcribed spacer in a tRNA transcript acts as a sponge for small RNAs to prevent transcriptional noise. Mol. Cell 58, 393–405.

42. Lalaouna, D., Prévost, K., Eyraud, A., and Massé, E. (2017). Identification of unknown RNA partners using MAPS. Methods San Diego Calif 117, 28–34.

43. Landt, S.G., Abeliuk, E., McGrath, P.T., Lesley, J.A., McAdams, H.H., and Shapiro, L. (2008). Small non-coding RNAs in Caulobacter crescentus. Mol. Microbiol. 68, 600–614.

44. Landt, S.G., Lesley, J.A., Britos, L., and Shapiro, L. (2010). CrfA, a small noncoding RNA regulator of adaptation to carbon starvation in Caulobacter crescentus. J. Bacteriol. 192, 4763–4775.

45. Langmead, B., Salzberg, S.L., and Slazberg, S.L. (2012). Fast gapped-read alignment with Bowtie 2. Nat. Methods 9, 357–359.

46. Laub, M.T., Chen, S.L., Shapiro, L., and McAdams, H.H. (2002). Genes directly controlled by CtrA, a master regulator of the Caulobacter cell cycle. Proc. Natl. Acad. Sci. U. S. A. 99, 4632–4637.

47. Levine, E., Zhang, Z., Kuhlman, T., and Hwa, T. (2007). Quantitative characteristics of gene regulation by small RNA. PLoS Biol. 5, e229.

48. Li, H., Handsaker, B., Wysoker, A., Fennell, T., Ruan, J., Homer, N., Marth, G., Abecasis, G., and Durbin, R. (2009). The Sequence Alignment/Map format and SAMtools. Bioinformatics 25, 2078–2079.

49. Liu, D., Chang, X., Liu, Z., Chen, L., and Wang, R. (2011). Bistability and oscillations in gene regulation mediated by small noncoding RNAs. PloS One 6, e17029.

50. Lott, S.C., Schäfer, R.A., Mann, M., Backofen, R., Hess, W.R., Voβ, B., and Georg, J. (2018a). GLASSgo – Automated and Reliable Detection of sRNA Homologs From a Single Input Sequence. Front. Genet. 9.

51. Lott, S.C., Schäfer, R.A., Mann, M., Backofen, R., Hess, W.R., Voβ, B., and Georg, J. (2018b). GLASSgo – Automated and Reliable Detection of sRNA Homologs From a Single Input Sequence. Front. Genet. 9.

52. Love, M.I., Huber, W., and Anders, S. (2014). Moderated estimation of fold change and dispersion for RNA-seq data with DESeq2. Genome Biol. 15, 550.

53. Madeira, F., Park, Y. mi, Lee, J., Buso, N., Gur, T., Madhusoodanan, N., Basutkar, P., Tivey, A.R.N., Potter, S.C., Finn, R.D., et al. (2019). The EMBL-EBI search and sequence analysis tools APIs in 2019. Nucleic Acids Res. 47, W636–W641.

54. Mandin, P., and Guillier, M. (2013). Expanding control in bacteria: interplay between small RNAs and transcriptional regulators to control gene expression. Curr. Opin. Microbiol. 16, 125–132.

55. Marczynski, G.T., and Shapiro, L. (2002). Control of chromosome replication in caulobacter crescentus. Annu. Rev. Microbiol. 56, 625–656.

56. Marczynski, G.T., Lentine, K., and Shapiro, L. (1995). A developmentally regulated chromosomal origin of replication uses essential transcription elements. Genes Dev. 9, 1543–1557.

57. Markham, N.R., and Zuker, M. (2005). DINAMelt web server for nucleic acid melting prediction. Nucleic Acids Res. 33, W577–581.

58. Marks, M.E., Castro-Rojas, C.M., Teiling, C., Du, L., Kapatral, V., Walunas, T.L., and Crosson, S. (2010). The genetic basis of laboratory adaptation in Caulobacter crescentus. J. Bacteriol. 192, 3678–3688.

59. McClure, R., Balasubramanian, D., Sun, Y., Bobrovskyy, M., Sumby, P., Genco, C.A., Vanderpool, C.K., and Tjaden, B. (2013). Computational analysis of bacterial RNA-Seq data. Nucleic Acids Res. 41, e140.

60. McGrath, P.T., Iniesta, A.A., Ryan, K.R., Shapiro, L., and McAdams, H.H. (2006). A dynamically localized protease complex and a polar specificity factor control a cell cycle master regulator. Cell 124, 535–547.

61. Mitarai, N., Benjamin, J.-A.M., Krishna, S., Semsey, S., Csiszovszki, Z., Massé, E., and Sneppen, K. (2009). Dynamic features of gene expression control by small regulatory RNAs. Proc. Natl. Acad. Sci. U. S. A. 106, 10655–10659.

62. Mohapatra, S.S., Fioravanti, A., Vandame, P., Spriet, C., Pini, F., Bompard, C., Blossey, R., Valette, O., and Biondi, E.G. (2020). Methylation-dependent transcriptional regulation of crescentin gene (creS) by GcrA in Caulobacter crescentus. Mol. Microbiol.

63. Murray, S.M., Panis, G., Fumeaux, C., Viollier, P.H., and Howard, M. (2013). Computational and genetic reduction of a cell cycle to its simplest, primordial components. PLoS Biol. 11, e1001749.

64. Nitzan, M., Rehani, R., and Margalit, H. (2017). Integration of Bacterial Small RNAs in Regulatory Networks. Annu. Rev. Biophys. 46, 131–148.

65. Panis, G., Lambert, C., and Viollier, P.H. (2012). Complete genome sequence of Caulobacter crescentus bacteriophage φCbK. J. Virol. 86, 10234–10235.

66. Panis, G., Murray, S.R., and Viollier, P.H. (2015). Versatility of global transcriptional regulators in alpha-Proteobacteria: from essential cell cycle control to ancillary functions. FEMS Microbiol. Rev. 39, 120–133.

67. Pini, F., Frage, B., Ferri, L., De Nisco, N.J., Mohapatra, S.S., Taddei, L., Fioravanti, A., Dewitte, F., Galardini, M., Brilli, M., et al. (2013). The DivJ, CbrA and PleC system controls DivK phosphorylation and symbiosis in Sinorhizobium meliloti. Mol. Microbiol. 90, 54–71.

68. Pini, F., De Nisco, N.J., Ferri, L., Penterman, J., Fioravanti, A., Brilli, M., Mengoni, A., Bazzicalupo, M., Viollier, P.H., Walker, G.C., et al. (2015). Cell Cycle Control by the Master Regulator CtrA in Sinorhizobium meliloti. PLoS Genet. 11, e1005232.

69. Quon, K.C., Marczynski, G.T., and Shapiro, L. (1996). Cell cycle control by an essential bacterial two-component signal transduction protein. Cell 84, 83–93.

70. Quon, K.C., Yang, B., Domian, I.J., Shapiro, L., and Marczynski, G.T. (1998). Negative control of bacterial DNA replication by a cell cycle regulatory protein that binds at the chromosome origin. Proc. Natl. Acad. Sci. U. S. A. 95, 120–125.

71. Ramírez, F., Dündar, F., Diehl, S., Grüning, B.A., and Manke, T. (2014). deepTools: a flexible platform for exploring deep-sequencing data. Nucleic Acids Res. 42, W187–W191.

72. Reisenauer, A., and Shapiro, L. (2002). DNA methylation affects the cell cycle transcription of the CtrA global regulator in Caulobacter. EMBO J. 21, 4969–4977.

73. Robinson, J.T., Thorvaldsdóttir, H., Winckler, W., Guttman, M., Lander, E.S., Getz, G., and Mesirov, J.P. (2011). Integrative genomics viewer. Nat. Biotechnol. 29, 24–26.

74. Ryan, K.R., Huntwork, S., and Shapiro, L. (2004). Recruitment of a cytoplasmic response regulator to the cell pole is linked to its cell cycle-regulated proteolysis. Proc. Natl. Acad. Sci. U. S. A. 101, 7415–7420.

75. Schneider, C.A., Rasband, W.S., and Eliceiri, K.W. (2012). NIH Image to ImageJ: 25 years of image analysis. Nat. Methods 9, 671–675.

76. Schrader, J.M., Zhou, B., Li, G.-W., Lasker, K., Childers, W.S., Williams, B., Long, T., Crosson, S., McAdams, H.H., Weissman, J.S., et al. (2014). The coding and noncoding architecture of the Caulobacter crescentus genome. PLoS Genet. 10, e1004463.

77. Siam, R., Brassinga, A.K.C., and Marczynski, G.T. (2003). A dual binding site for integration host factor and the response regulator CtrA inside the Caulobacter crescentus replication origin. J. Bacteriol. 185, 5563–5572.

78. Sievers, F., Wilm, A., Dineen, D., Gibson, T.J., Karplus, K., Li, W., Lopez, R., McWilliam, H., Remmert, M., Söding, J., et al. (2011). Fast, scalable generation of high-quality protein multiple sequence alignments using Clustal Omega. Mol. Syst. Biol. 7, 539.

79. Skerker, J.M., and Laub, M.T. (2004). Cell-cycle progression and the generation of asymmetry in Caulobacter crescentus. Nat. Rev. Microbiol. 2, 325–337.

80. Skerker, J.M., Prasol, M.S., Perchuk, B.S., Biondi, E.G., and Laub, M.T. (2005). Two-component signal transduction pathways regulating growth and cell cycle progression in a bacterium: a system-level analysis. PLoS Biol. 3, e334.

81. Sommer, J.M., and Newton, A. (1988). Sequential regulation of developmental events during polar morphogenesis in Caulobacter crescentus: assembly of pili on swarmer cells requires cell separation. J. Bacteriol. 170, 409–415.

82. Taylor, J.A., Ouimet, M.-C., Wargachuk, R., and Marczynski, G.T. (2011). The Caulobacter crescentus chromosome replication origin evolved two classes of weak DnaA binding sites. Mol. Microbiol. 82, 312–326.

83. Tien, M., Fiebig, A., and Crosson, S. (2018). Gene network analysis identifies a central post-transcriptional regulator of cellular stress survival. ELife 7.

84. Tjaden, B. (2015). De novo assembly of bacterial transcriptomes from RNA-seq data. Genome Biol. 16, 1.

85. Wurihan, W., Wunier, W., Li, H., Fan, L.F., and Morigen, M. (2016). Trans-translation ensures timely initiation of DNA replication and DnaA synthesis in Escherichia coli. Genet. Mol. Res. GMR 15.

86. Zhan, Y., Yan, Y., Deng, Z., Chen, M., Lu, W., Lu, C., Shang, L., Yang, Z., Zhang, W., Wang, W., et al. (2016). The novel regulatory ncRNA, NfiS, optimizes nitrogen fixation via base pairing with the nitrogenase gene nifK mRNA in Pseudomonas stutzeri A1501. Proc. Natl. Acad. Sci. U. S. A. 113, E4348–4356.

87. Zhou, B., Schrader, J.M., Kalogeraki, V.S., Abeliuk, E., Dinh, C.B., Pham, J.Q., Cui, Z.Z., Dill, D.L., McAdams, H.H., and Shapiro, L. (2015). The global regulatory architecture of transcription during the Caulobacter cell cycle. PLoS Genet. 11, e1004831.

88. Zuker, M. (2003). Mfold web server for nucleic acid folding and hybridization prediction. Nucleic Acids Res. 31, 3406–3415.

